# Pallidin function in drosophila surface glia regulates sleep and is dependent on amino acid availability

**DOI:** 10.1101/2022.05.03.490434

**Authors:** Hui Li, Sami Aboudhiaf, Sandrine Parrot, Céline Scote-Blachon, Claire Benetollo, Jian-Sheng Lin, Laurent Seugnet

## Abstract

The Pallidin protein is a component of a multimeric complex named the Biogenesis of Lysosome-related Organelles Complex 1 (BLOC1) that regulates specific endosomal function and transmembrane protein trafficking in many different cell types. In the brain, defective BLOC1 function has been linked to schizophrenia, a neuropsychiatric disorder with highly prevalent sleep disruptions, and to impaired cognitive abilities in healthy subjects. In animal models, defective BLOC1 function also impairs behavior, memory, neurotransmission systems and metabolism. This growing body of experimental evidence suggest an involvement of BLOC1 in sleep/wake regulation. Here, we used *Drosophila* molecular genetics and conditional, cell-type specific knockdown strategy to address this question. We show that down-regulation of a central subunit of BLOC1, Pallidin, in the surface glia, the *Drosophila* equivalent of the blood brain barrier, is sufficient to reduce, fragment and delay nighttime sleep at the adult stage and in a circadian clock dependent manner. Other members of the BLOC1 complex appear to be involved in this surface glia-dependent sleep regulation. In agreement with a BLOC1 involvement in amino acid transport, down-regulation of the Large neutral Amino acid Transporter 1 (LAT1)-like transporters *JhI-21* and minidiscs, phenocopy the down-regulation of *pallidin*. Similar results were obtained by inhibiting the TOR amino acid signaling pathway. Supplementing food with essential amino acids normalizes the sleep/wake phenotypes of *pallidin* and *JhI-21* down-regulation. Furthermore, we identify a role for *pallidin* in the subcellular trafficking of *JhI-21* in surface glial cells. Finally, we provide evidence that *Pallidin* function in surface glia is required for GABAergic neurons activation involved in promoting sleep. Taken together, these data identify a novel role for BLOC1 that, through LAT1-like transporters subcellular trafficking modulates essential amino acid availability and GABAergic sleep/wake regulation.

## Introduction

Identified over 20 years ago, the Biogenesis of Lysosome-related Organelles Complex 1 (BLOC1) is an octameric complex linked to endosomal compartments and the cytoskeleton^1–3^. The genes coding for the 8 subunits in mice: pallidin, dysbindin, BLOS1, BLOS2, BLOS3, cappuccino, muted and snapin are broadly expressed within the brain and in peripheral tissues. The complex regulates the trafficking of various receptors and transporters notably in melanocytes, but also in neurons, and appears to play a prominent role in the biogenesis of recycling endosomes^4–7^. Protein-protein interactions within the complex contribute to its stability such that lack of any one subunit leads to dramatically reduced protein levels of the other subunits^8–17^. Unexpectedly, mice bearing severe or complete loss of function mutations in BLOC1 genes are viable and fertile^1, 18^. They display common phenotypes, originating from defects in highly specialized lysosome-related organelles, such as reduced pigmentation due to impaired retinal and epidermal melanosomes, or extended bleeding times resulting from the lack of dense granules in platelets. In humans, mutations in BLOC1 genes and other functionally related genes are found in the Hermansky-Pudlack syndrome, which is similarly characterized by albinism and increased bleeding time^19^. In addition, genetic studies have identified variants of the dysbindin gene and other BLOC1 genes as risk factors for developing schizophrenia. While the latter results are debated^20^, several postmortem studies have reported reduced levels of dysbindin mRNA and protein in the brain of schizophrenics^1, 3, 21–23^ providing additional support for the link. Furthermore, genome-wide association studies also reported a link between dysbindin genetic variants and cognitive abilities including memory encoding and executive function^24, 25^. These findings have led to investigations aiming at deciphering the role of BLOC1 in neuronal function^1^ using not only mouse models defective for individual BLOC1 gene function but also the *Drosophila* model, in which the complex is well conserved^9, 26–30^. These studies confirmed the involvement of BLOC1 in behavior and memory. They identified potential cellular and molecular mechanisms such as abnormal glutamatergic, GABAergic and dopaminergic transmission^1, 9, 26, 27, 29, 31–33^ but also changes in metabolism and amino acid transport in the brain^34, 35^. However, the vast majority of these studies have been carried out in animals with spontaneous or artificial mutations, thus bearing constitutive and ubiquitous alteration in gene function. This approach makes the interpretation of the resulting phenotypes challenging, given the broad expression of BLOC1 in many cell types. Furthermore, the complex is expressed during development, where it can regulate neurite outgrowth and dendritic spine formation, and be involved in neurodevelopmental diseases such as autism spectrum disorders^1, 20, 36, 37^.

In this context, and apart from one recent report^32^, the involvement of BLOC1 genes in sleep-wake regulation has not been investigated. Further investigation is required given the critical implication of sleep in memory, in the regulation of neurotransmission, and in neuropsychiatric disorders, notably schizophrenia. The highly prevalent sleep disruption in schizophrenia could be in part a consequence of the pathology^38–40^, and is also an aggravating factor since sleep deprivation has been shown to induce schizophrenia-like behavior in human^41^ and in rodent models^42^. Here we investigated the potential role of the BLOC1 complex in sleep/wake regulation in the *Drosophila model*, using a conditional knockdown of one major component of the BLOC1 complex, *pallidin*. The pool of Pallidin in the cell is almost entirely associated with the complex and this protein plays a central role through its interactions with dysbindin, BLOS1, and cappuccino^2, 28^, its loss leading to their degradation^1, 36^. Using that genetic strategy, we focused our study on glial cells, since they play a major role in both metabolism and the modulation of neurotransmission.

## Materials and methods

### Fly stocks and husbandry

Flies were raised on standard food containing inactivated yeast, cornmeal, agar, molasses, sucrose and maintained at 25°C, 60% humidity in a 12hr: 12hr Light: Dark (LD) cycle, except otherwise mentioned. The following stocks obtained were used in this study: from the Bloomington Drosophila Stock Center: *Canton S, UAS-dicer2, UAS-Pallidin-RNAi^HMS05728^, repo-Gal4; GMR54C07-Gal4, GMR85G01-Gal4, Tub-Gal80^ts^, UAS-snapin-RNAi^JF02692^, UAS-JhI-21-RNAi^HMS02271^, UAS-Raptor-RNAi^HMS02306^, UAS-TOR-RNAi^HMS00904^, Df(3L)BSC675, UAS-nuclear-lacZ*, vGAT-Gal4; from the Vienna Drosophila Resource Center: *<u>UAS-pallidin-RNAi</u>^GD13391^, UAS-Blos2-RNAi^GD34355^,UAS-JhI-21-RNAi^KK112996^, UAS-Mnd-RNAi^KK102686^, UAS-Mnd-RNAi^GD453^, UAS-RNAi-CD98Hc^GD46622^, UAS-RNAi-CD98Hc^HMC04939^*; from the National Institute of Genetics, Mishima: *UAS-Blos1-RNAi^CG30077R-3^, UAS-dysbindin-RNAi^CG6856R-1^*; from A. Borst, Max Planck Institute of neurobiology: *VGAT-lexA*.

From Y. Aso, Janelia farm, HHMI: *TH-lexA*; from S. Birman, Centre National de La recherche Scientifique: *LexAOP-TrpA1, DAT^fumin^*; from I. Alliaga London Institute of Medical Sciences: *Ilp2-Gal4*; from B. Mollereau, Ecole Normale Supérieure de Lyon: *UAS-GFP, UAS-mCherry-GFP-LC3*; from D. Dickman, University of Southern California: *pallidin ^Δ1^*, from C Klämbt, Munster University: *UAS-CD4-spGFP1-10, LexAOP-CD4-spGFP11*/TM6. All lines were outcrossed 3 times to a Canton S reference strain or to other lines previously outcrossed to Canton S before being used for experimentation. For standard Gal4-UAS experiments, the Gal4 and UAS parental lines outcrossed to Canton S served as genetic background controls. In most experiments, the UAS-dicer2 transgene was added to the genetic background to increase the efficiency of the knockdown. For the experiments involving the TARGET system^43^ or *LexAOP-TrpA1* expression, flies were raised at 18°C to prevent Gal4 and TrpA1 activity, respectively. Evaluation of *pallidin* transcripts by QPCR shows that *UAS-Pallidin-RNAi^HMS05728^* combined with *da-Gal4* significantly downregulates the *pallidin* transcripts in the whole body (Supplemental data, Figure S1).

### Sleep recording, sleep deprivation and circadian rhythms evaluation

Freshly hatched female flies were collected under CO_2_ anesthesia and loaded individually at age 2-5 days into 5x65mm long glass tubes containing standard food medium. Sleep was recorded in a Light: Dark 12h:12h (LD) cycle at 25°C, 60% humidity using the Trikinetics DAMS system (www.trikinetics.com), unless otherwise mentioned. Sleep parameters were evaluated across 3-6 days of baseline as described previously using 5 minutes immobility as criteria^44,45^. We used the Drosophila ARousal Tracking (DART) system^46^ for the evaluation of sleep intensity. Sleep experiments were repeated at least 3 times. Distribution and homogeneity as well as statistical group comparisons were evaluated using the Microsoft Excel plugin software Statel. Mean ± SEM is plotted and the p value shown is the highest obtained among post hoc comparisons. Black box areas indicate the scotophase in the 24h hourly sleep graphs. Sleep deprivation was carried out using a Sleep Nullifying Apparatus system (SNAP) as previously described^44, 47^, for 24hr starting at ZT0. For the evaluation of circadian rhythms, flies were recorded in constant darkness for 6 days, and the free-running period was calculated using locomotor data and the χ^2^ method in ActogramJ^48^.

### Immunofluorescence

Flies were anesthetized on ice and brains dissected quickly in cold phosphate-buffered saline (PBS) solution before being fixed for 25min in a 4% paraformaldehyde PBS solution at room temperature. A 5% normal goat serum (Sigma) in PBS with 0,3% TritonX was used for blocking during at least 45min, before overnight incubation at 4°C in 0,3% TritonX PBS with the primary antibody: rabbit anti-JhI-21 (1/250; kindly provided by Y. Grosjean, University of Burgundy, France^16, 26^), combined with mouse anti-GFP (8H11, from University of Iowa Developmental Studies Hybridoma Bank, 1/200), or mouse anti-β-galactosidase (Z378A from Promega, 1/1000). Secondary antibodies (anti-rabbit IgG-Alexa Fluor 633 and anti-mouse IgG-Alexa Fluor 488) were purchased from Thermo Fisher Scientific and used at 1/1000. Brains were mounted in Vectashield medium containing DAPI (Vector Laboratories). Fluorescence was observed using a confocal microscope (Zeiss LSM 800) with a 40x objective and Image J (FIJI) was used for image processing. DAPI fluorescence and β-galactosidase immunofluorescence were used to delineate the nuclei in neurons and surface glia respectively. For the quantification of anti-JhI-21 immunofluorescence in neuronal nuclei, the average pixel intensity in control brains was used to normalize the signal. For the quantification of anti-JhI-21 immunofluorescence in perineurial and subperineurial cells, the average pixel intensity of 5 neuronal nuclei was substracted to the signal detected in surface glial cells in the same optical section. The average pixel intensity in control subperineurial cells was the lowest observed in any cells and arbitrarily set to zero.

### GRASP experiments

The *vGAT-LexA* transgene was used to drive expression of *LexAOP-CD4-sp-GFP*^11^ in all GABAergic neurons, while *SG-Gal4* was used to drive *UAS-CD4-sp-GFP*^1–10^ in surface glia. Flies were dissected in PBS, fixed for 5min in a 4% paraformaldehyde PBS solution at room temperature, incubated for 10 min in 0,3% TritonX PBS containing 1μg/mL DAPI and 1:100 Phalloidin-rhodamine (*Abcam* ab235138), rinsed and mounted in PBS solution. Fluorescence was observed using a confocal microscope (Zeiss LSM 800) with a 40x objective and Image J (FIJI) was used for image processing.

### Leucine supplementation

Flies in Trikinetics tubes were recorded for 2 days and transferred at ZT0 from standard food to food containing either Leucine, valine or Tryptophan at 50mM or to tubes containing fresh standard food (control condition).

### L-DOPA feeding

For L-DOPA feeding experiments, flies in Trikinetics tubes were transferred at ZT0 from standard food to vehicle (1% agar 1% sucrose) for 24h (day 1). At ZT0 the following day, flies of each genotype were divided into two groups, one placed on fresh vehicle for 24h (“vehicle” condition) and the other one on vehicle plus L-DOPA dissolved at 3mg/ml (day 2). The % of nighttime sleep change: (day 2 sleep – day 1 sleep)/ day 1 sleep is displayed as a box plot. Food intake was evaluated using the CAFE assay^27^: in vials containing 5 flies, food was provided by a 5 μL calibrated capillary filled with liquid food containing 5 % sucrose, 5 % yeast extract, 1 % blue food coloring die and L-DOPA (3mg/ml). Experimental readings and capillaries replacement were carried out every 12 hours. An identical CAFE chamber without flies was used to determine evaporative losses (typically <10% of ingested volumes), which were subtracted from experimental readings.

### Analysis of tissue amino acid content using HPLC-LIFD

Single brain samples from drosophila were quickly dissected in 1 µL of Ringer’s solution and frozen at -80°C until tissue content analysis. The extraction was made similarly as for dopamine and serotonin content analysis, but with 3 µL of perchloric acid/EDTA/sodium bisulfite solution. Aspartate, glutamate, and GABA levels were quantified using HPLC with laser induced fluorescence detector (HPLC-LIFD) from adapted chromatographic conditions^49^ in a low-pressure binary gradient mode. The HPLC system (Shimadzu, Japan) consisted of a Prominence degasser, a high-pressure LC-30 AD pump (as pump A), a low-pressure LC-20 pump (as pump B), a 20-µL mixer and an SIL-30AC autosampler. The mixing of the mobile phases from both pumps and fluorescence detection using a ZETALIF laser induced fluorescence detector (Picometrics, France) were performed at 23-24°C before injection and after separation, respectively. The detection was carried out at the outlet of the column in a fused-silica capillary serving as detection cell (i.d. 75 µm, OD 375 µm, stripped out of its polyimide layer at 50 cm (effective length), total length 60 cm, towards waste). Excitation was performed using a 410-nm diode laser (Melles Griot, USA) and the emission wavelength was set at 490 nm. Separations were performed using a 100 × 2.1 mm Kinetex C18 core-shell 2.6 µm column equipped with a KrudKatcher Ultra HPLC in-line filter. The column oven was set to 45°C. 1 µL of each extract sample was diluted 100-fold in a Ringer’s solution. The derivatization reagents were added off-line in the 100 µL of the diluted extracts, by adding 20 µL of borate/NaCN solution (mixing solution [250:50, v/v] of 500 mmol/L borate buffer pH 8.7, and 87 mmol/L NaCN in water), 10µL of ultrapure water, and, in the end, 10 µL of NDA solution (2.925 mmol/L in acetonitrile/water, 50:50, v/v) for a 15-min reaction at 23-24°C^30^. 10 µL of the derivatized samples were placed in the autosampler and kept at 4°C before injection. The injection volume was 2 µL. Separations were performed at a flow rate of 0.3 mL/min (∼ 260 bars). The gradient elution conditions were as follows: buffer: 50 mmol/L phosphate buffer pH 6.8; mobile phase A: buffer/tetrahydrofuran [97:3 v:v]; mobile phase B: buffer/methanol/acetonitrile [35:10:55, v:v:v]; gradient: 22% B increased to 27 % B from 0.01 to 5.08 min, increased to 33 % B from 5.08 min to 6.41 min, increased to 41 % B from 6.41 to 14.52 min, increased to 100 % B from 14.52 to 15.85 min, kept at 100 % until 21.19 min, then decreased to 22 % B from 21.19 min to 22.52 min; kept at 22 % B until the end of the run. The acquisition time was 27.5 min. Chromatograms were acquired at a rate of 10 Hz (Lab Solutions software). The retention times were ∼1.8 min, ∼2.5 min, and ∼11.2 min for Asp, Glu, and GABA, respectively. Concentrations of the analytes in the extracts were determined by comparison of chromatographic peak areas with calibration curves derived from a mixture of the 3 synthetic standards (two points: 10^−6^, 5 × 10^−6^ mol/L).

### Analysis of dopamine and serotonin content using capillary HPLC

5 brains were quickly dissected from cold-anesthetized flies and frozen in 5 μL of ringer solution at -80°C until being processed for tissue content analysis. More precisely, 15 µL of ice-cold 0.1 mol/L perchloric acid containing 1.34 mmol/L EDTA and 0.05%, w/v sodium bisulfite were added to each sample and samples were then sonicated in an ultra-sonic bath for 12 min. The homogenates were centrifuged at 15,000 × g for 20 min at +4°C, and the supernatants analyzed for monoamine content on the same day or the following day when a dilution of sample was required due to some very high dopamine concentrations (saturated peaks). The HPLC system consisted of a Prominence degasser, an LC-30 AD pump, and an SIL-30AC autosampler from Shimadzu. Detection was carried out at 43°C using a Decade II electrochemical detector fitted with a 0.7 mm glass carbon working electrode, an Ag/AgCl reference electrode, and a 25-µm spacer (cell volume 80 nL, Antec). Separations were performed in the detector oven at 43°C using a capillary 150 × 0.5 mm Zorbax-SB C18 5 µm column. The mobile phase, which was pumped at a flow rate of 10 µL/min, consisted of 100 mmol/L sodium phosphate containing 0.1 mmol/L EDTA, 2 mmol/L octane sulfonate, and 0.01% triethylamine, with a pH adjusted to 5.5 with sodium hydroxide, filtered through a 0.22 µm membrane before use and mixed with 17% methanol (v:v). Analytes were detected at an oxidation potential of 700 mV versus the reference electrode. Chromatograms were acquired at a rate of 10 Hz using Lab Solutions software. The acquisition time was 40 min per sample. The injection volume was 1 µL. Concentrations of dopamine and serotonin in the extracts were determined by comparison of chromatographic peak areas with calibration curves derived from a mixture of synthetic standards containing dopamine and serotonin (two points: 2.5 × 10^−7^ and 5 × 10^−7^ mol/L). The average concentration in the group on vehicle was used to normalize the results which are expressed as percentage of vehicle controls.

### Quantitative real time PCR

Total RNA was extracted from 10 homogenized bodies using the RNeasy Minikit with DNase treatment (Qiagen) and reverse transcription was performed using SuperScript IV Reverse Transcriptase (Invitrogen). qPCR amplifications were performed with the Rotor-Gene SYBR Green PCR Kit (Qiagen), on a Rotor-Gene Qcycler (Qiagen). Absolute quantification of cDNA copy number was obtained by relating the Ct value to a standard curve made with serial dilutions of a reference DNA. The copy numbers obtained for the *Ribosomal Protein 49 (RP49)* gene were used as controls. The values shown correspond to the copy number of *pallidin* divided by the copy number for RP49 amplified transcripts. Primers were designed with Primer-Blast software (National Centre for Biotechnology Information/NCBI, Bethesda, USA) and purchased from Eurogentec.

## Results

### Pallidin is required in adult Drosophila surface glia for normal sleep-wake regulation in a circadian clock dependent manner

Given that pallidin is part of the evolutionary conserved BLOC1 complex^28^, we used the *Drosophila* model and the Gal4-UAS system to downregulate *pallidin* with *UAS-pallidin-RNAi* transgenes in different cell types^50^ to investigate its functional implication in sleep-wake regulation. Inhibition of *pallidin* expression in all neurons as well as in all glia significantly affected sleep/wake (Figure 1, see Table 1 for all the genotypes used in this study). We focused our analysis on glia, given that the role of *pallidin* in those cells has not been explored, despite their significant role in metabolism, nutriment transport, trafficking and ubiquitous impact on all brain cells. Inhibition in all glial cells resulted in a significant reduction of night sleep especially in the first half of the dark period, the primary sleep period in *Drosophila* (Figure 1A)^44, 51^. Specific Gal4 drivers were used to determine which glial cell type was responsible for the phenotype observed with pan-glial knockdown. As shown in Figure 1B-C, the inhibition of *pallidin* expression in perineurial or subperineurial cells, collectively called surface glia (SG), was sufficient to replicate the delayed and reduced night sleep phenotype obtained upon pan-glial inhibition, using two independent UAS-RNAi constructs (Supplemental Figure S1). The most consistently observed phenotype was a reduction in sleep amounts in the first half of the night, suggesting a difficulty initiating sleep at a time where flies display the highest amount of consolidated sleep^45^ and subperineurial glial cells their highest endocytic activity^52^. The reduction in average nighttime sleep bout duration was less consistent across drivers and RNAi constructs (Supplemental Figure S2-3) and for one of the *UAS-pallidin-RNAi* constructs, a sleep reduction and fragmentation during daytime was also observed (Figure S2B, S3B). Expression of a UAS-RNAi against *sugar baby*, a gene sharing sequences with *pallidin* and co-targeted by the *UAS-RNAi-Pallidin^<u>GD13391</u>^* construct, did not affect sleep (Supplemental Figure S1). Subperineurial cells are connected by septate junctions and cover the surface of the brain, controlling the exchanges with the circulating hemolymph^53^. Together with the perineurial cells, these cells constitute the invertebrate equivalent of the Blood-brain-barrier and are referred here as surface glia (SG). The *9-137-Gal4* driver line^53^, which is expressed in both perineurial and subperineurial glia, displayed a severe sleep loss phenotype during the nighttime as well as during the daytime. Although reported as SG specific^52, 53^, *9-137-Gal4* is also expressed at low levels in neurons of the *pars intercerebralis* (Supplemental Figure S4), which include Insulin-like-peptide 2 (ilp2) producing neurons, reported to play a role in sleep/wake regulation^54–57^. We combined *9-137-Gal4* with the *elav-Gal80* transgene to inhibit Gal4 activity in all neurons and restrict it to SG. This compound driver (SG-Gal4) showed no detectable expression in pars intercerebralis neurons and was still able to induce sleep loss with *UAS-pallidin-RNAi* (Supplemental Figure S4, Figure 1 C). In addition, driving *UAS-pallidin-RNAi* specifically in the ilp2 producing neurons with an *Ilp2-Gal4* driver did not affect sleep/wake (Supplemental Figure S5) further supporting the conclusion that the phenotype observed with 9-137-Gal4 is due to *pallidin* inhibition in surface glia.

**Figure 1:**
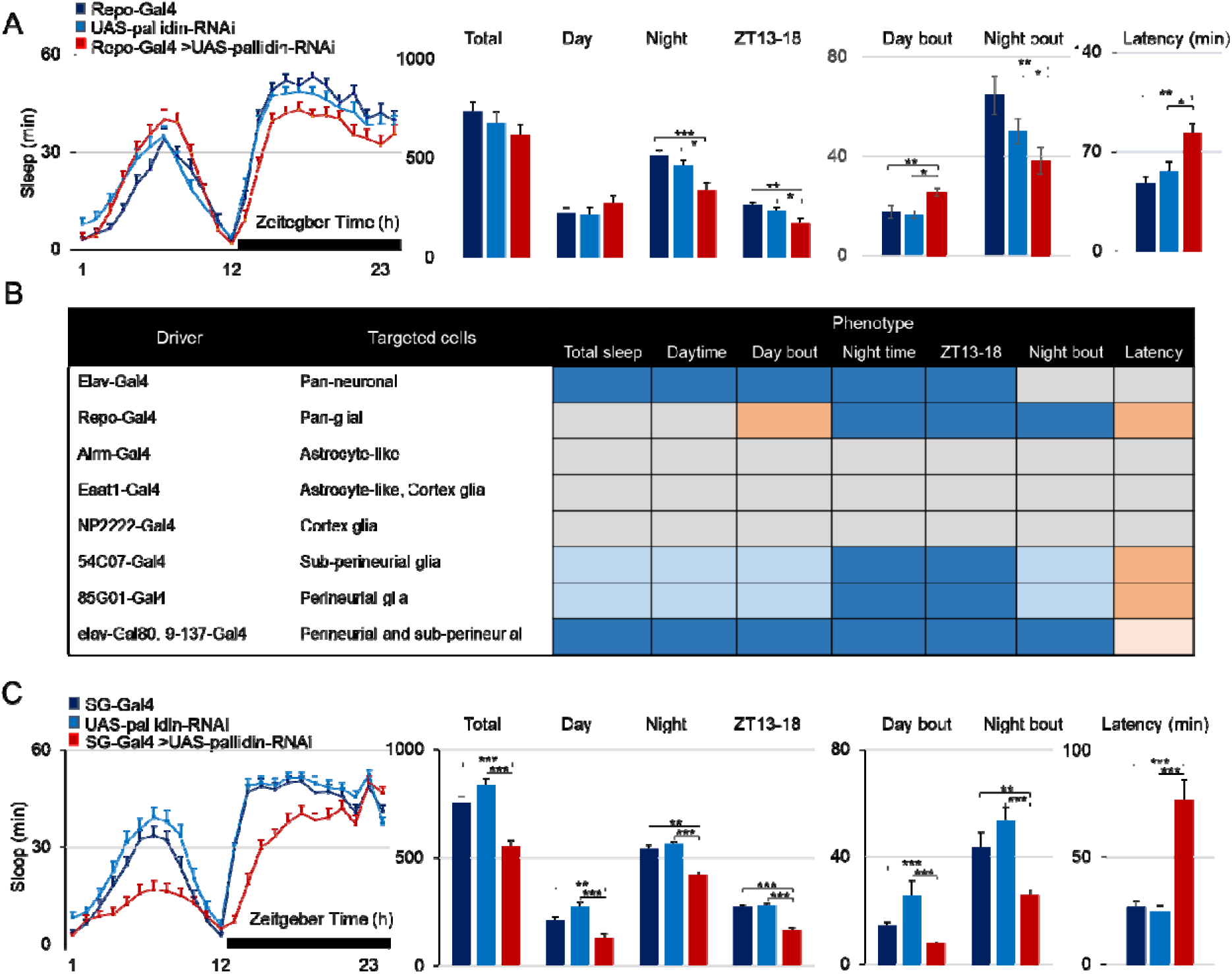
pallidin knockdown in surface glia results in delayed, reduced and fragmented night sleep. **A)** Baseline daily sleep of flies expressing a *UAS-pallidin-RNAi* driven in all glial cells (repo-Gal4 driver) in red, compared to the genetic control groups in blue. **B**) Comparison of sleep parameters upon expression of *UAS-pallidin-RNAi* under the control of several Gal4 drivers. 54C07-Gal4, 85G01-Gal4 and 9-137-Gal4 elav-Gal80 were crossed to both *UAS-pallidin-RNAi*^GD13391^ and *UAS-pallidin-RNAi*^HMS05728^, see supplemental data (Figure S1-S3). Dark blue color indicate significant increase and salmon color significant decrease in amount/duration of the parameters in the knockdown condition compared to both the Gal4 and UAS controls (p<0.05; Kruskal-Wallis with post hoc comparisons between control and knockdown conditions). Lighter color indicate that the changes were observed with the *UAS-pallidin-RNAi*^HMS05728^ only. **C**) Baseline daily sleep of flies expressing *UAS-pallidin-RNAi*, driven in subperineurial and perineurial cells (9-137-Gal4 combined with elav-Gal80 (SG) As in A, night sleep is reduced, especially during the second half of the night (ZT 13-18, middle left bar graph), as a consequence of prolonged night sleep latency (far right bar graph). Night sleep is also significantly fragmented (middle right bar graph). n=28-35 for each condition., *: p<0.05; **: p<0.005; ***: p<0.0005; Kruskal-Wallis with post hoc comparisons between control and knockdown conditions.

**Table 1.**
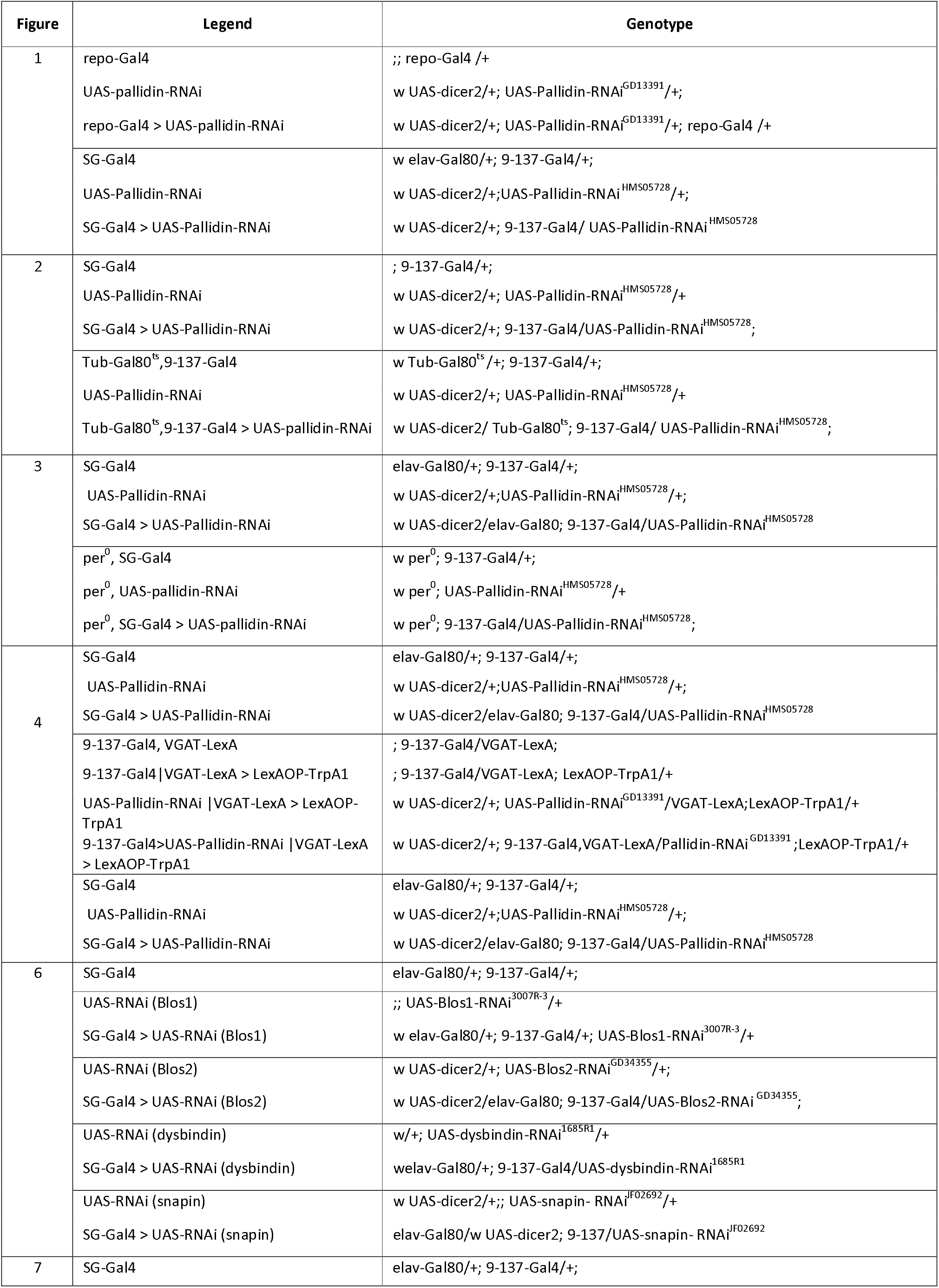
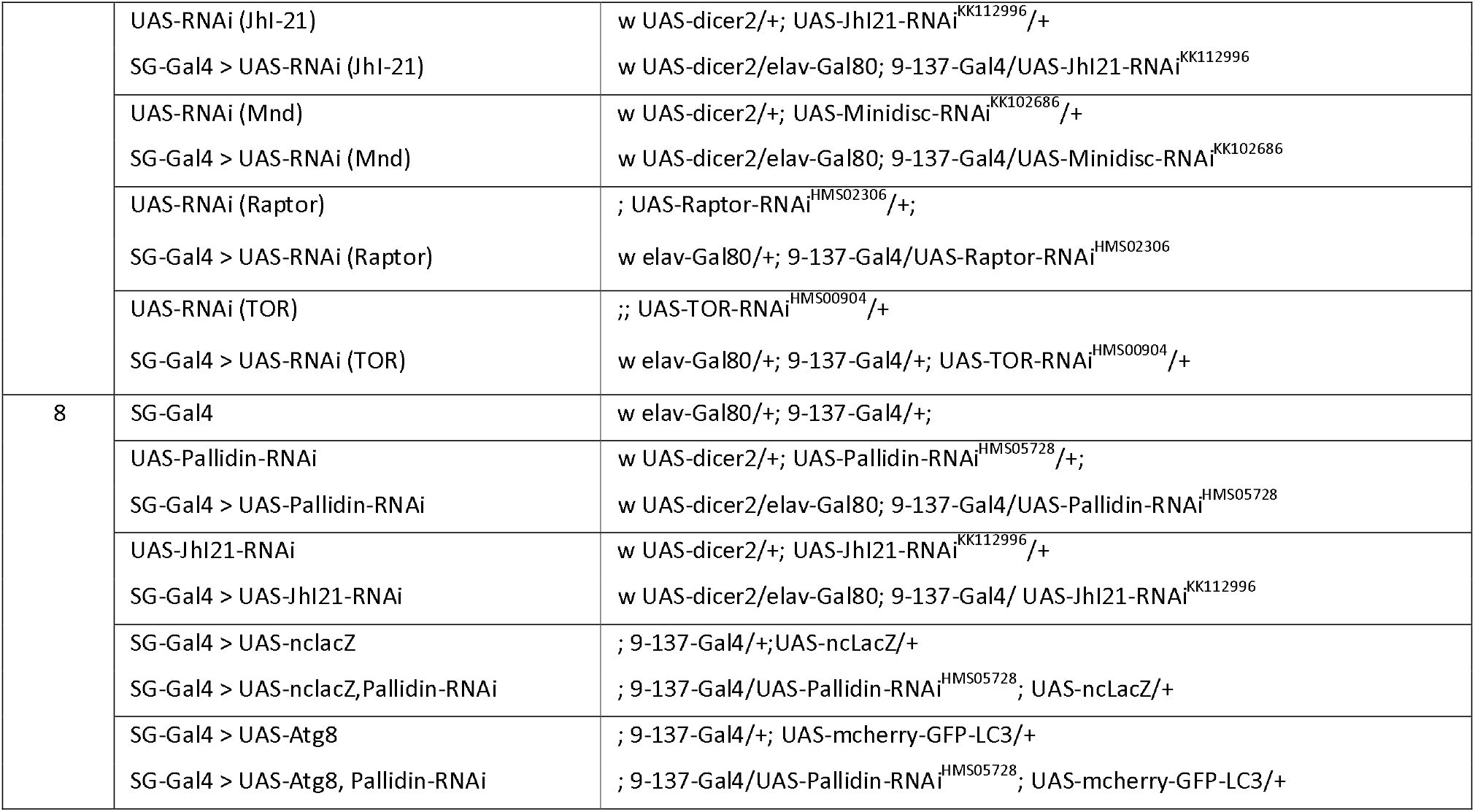

To determine whether *pallidin* is required for sleep regulation at the adult stage or earlier during development, we used the TARGET system to control the timing of *UAS-pallidin-RNAi* expression^43^. In flies raised and monitored at 18°C, *UAS-pallidin-RNAi* expression is inhibited by the thermosensitive Gal80^ts^ inhibitor of Gal4: in these conditions daytime sleep was still reduced in the experimental flies compared to controls, but no sleep reduction was observed during the night (Figure 2B). This latter lack of effect was not due to differences in sleep/wake regulation at 18°C, since nighttime sleep reduction was readily observed at 18°C in absence of Gal80^ts^ expression (Figure 2A). When the adult flies are transferred to 25°C, however, we observed reduced sleep amounts especially during the first half of the dark phase (Figure 2B). This phenotype was milder than in the absence of the *tub-Gal80*^ts^ transgene. Increasing the temperature to 30 or 32°C did not result in a more severe phenotype, which could be attributed to the fact that all flies show nocturnal behavior at that temperature, independently of circadian clock regulation^58^. In contrast, daytime sleep remained lower in the experimental flies compared to control, suggesting that this phenotype is independent from the *pallidin* knockdown. These results suggest that *pallidin* is required at the adult stage to regulate sleep/wake during the first half of the night. We therefore focused our investigations to this particular aspect of sleep-wake regulation.

**Figure 2:**
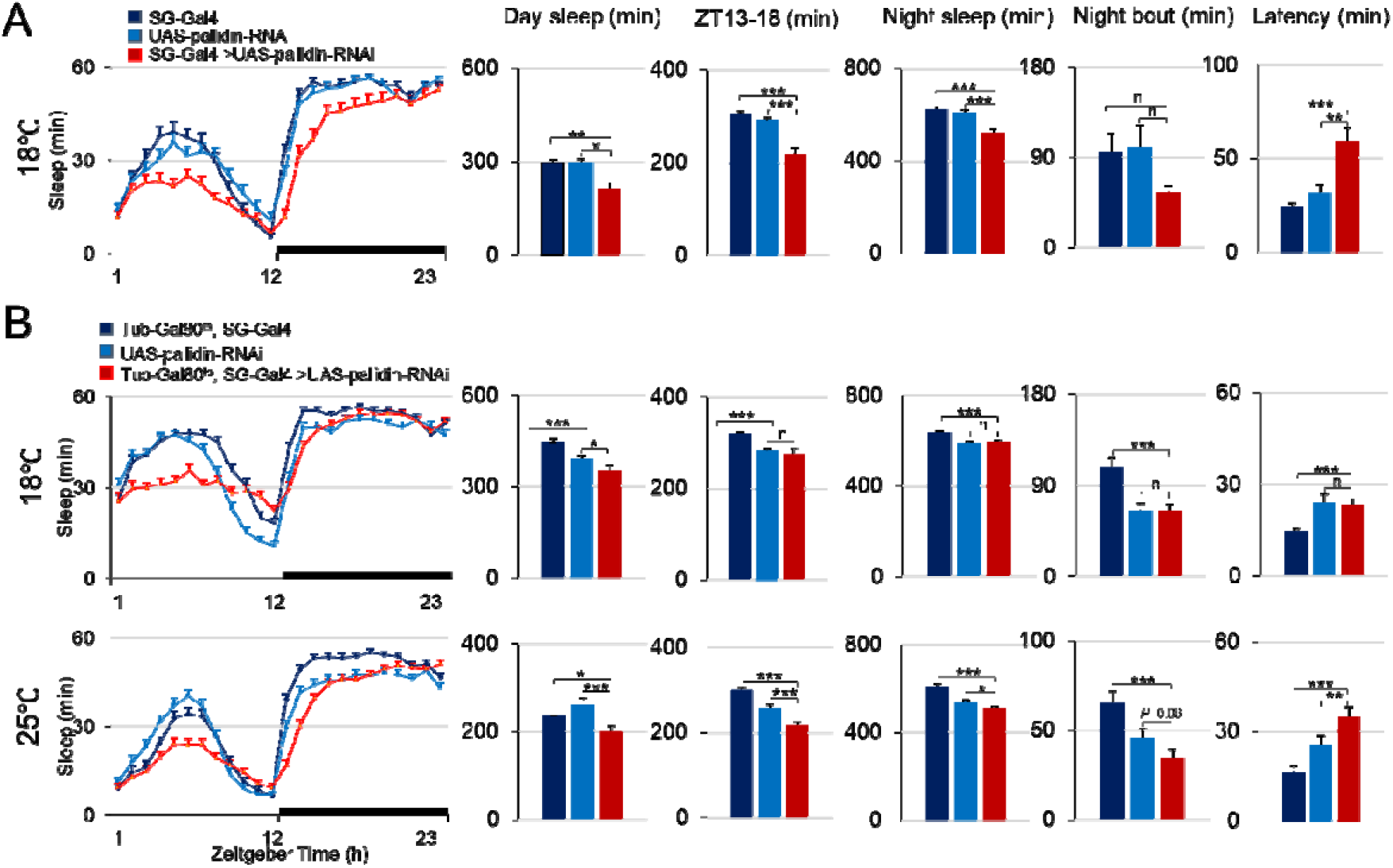
Pallidin downregulation in surface glia during adulthood is sufficient to impact sleep/wake regulation. A) pallidin knockdown flies (red) display delayed, reduced and fragmented sleep at 18°C, compared to controls (blue), similar to results obtained at 25°C. *: p<0.05, **: p<0.005, ***: p<0.0005; Kruskal-Wallis with post hoc comparisons between control and knockdown conditions N= 38-42 B) *Pallidin* knockdown was restricted to adulthood using the *Tub-Gal80* construct and allowing the flies to develop at 18°C. Nighttime sleep in *pallidin* knockdown flies is similar to the control groups at 18°C. At 25°C the night sleep of *pallidin* knockdown flies was reduced and delayed. Average night sleep bout duration was marginally reduced compared to controls. *: p<0.05, **: p<0.005, ***: p<0.0005; Kruskal-Wallis with post hoc comparisons between control and knockdown conditions, N= 84-93.

Sleep intensity is more pronounced during nighttime as assessed by sleep probing using sensory stimulation and video recording (^46, 59^, see methods). Interestingly, using this method, nighttime sleep intensity did not differ in *SG>pallidin-RNAi* flies, suggesting that the early night sleep loss is not originating from the inability of the flies to reach deep sleep stages (Supplemental Figure S6). The knockdown flies also showed normal localization close to the food during sleep (Supplemental Figure S6).

To identify the processes affected by *pallidin* knockdown, we assessed the two main inputs involved in sleep/wake regulation: sleep homeostasis and the circadian clock. *SG>pallidin-RNAi* flies displayed a sleep rebound in the first 6h following a 24h total sleep deprivation (Supplemental Figure S7), and longer day sleep bout duration compared to baseline, indicating that they were able to elicit a sleep homeostatic response. However, the percentage of sleep recovered was smaller than in the controls, indicating that the amplitude of the sleep homeostatic response is reduced in the *pallidin* knockdown flies. To assess the influence of light and of the circadian clock, flies were placed in constant darkness. In those conditions *SG>pallidin-RNAi* flies displayed a typical sleep/wake rhythm with reduced sleep during the presumptive light phase and a free-running period of 25.1±0.6 hr, comparable to 25±0.4 and 24.3±0.4 observed for the UAS and Gal4 controls respectively (N=30-32). As in Light: Dark conditions, *SG>pallidin-RNAi* displayed lower sleep amounts compared to the controls during the early part of the presumptive night (Figure 3 A-B), while the average sleep bout duration was not significantly reduced. These results indicate that the circadian clock is functional and that early night sleep loss in knockdown flies is independent from light exposure. To assess the implication of the circadian clock, the *per^0^* mutation, known to completely abolish the cycling of the molecular oscillator^60^, was introduced into the genetic background of the flies. The down-regulation of *pallidin* in SG was no longer disrupting sleep in presence of the per^0^ mutation, suggesting that the processes affected by *pallidin* are dependent on the circadian clock (Figure 3C-D). Altogether, these results provide strong evidence that *pallidin* is required in SG for sleep/wake regulation: the knockdown in SG results in sleep reduction and sleep fragmentation, especially during the early night phase, in a circadian clock dependent manner.

**Figure 3:**
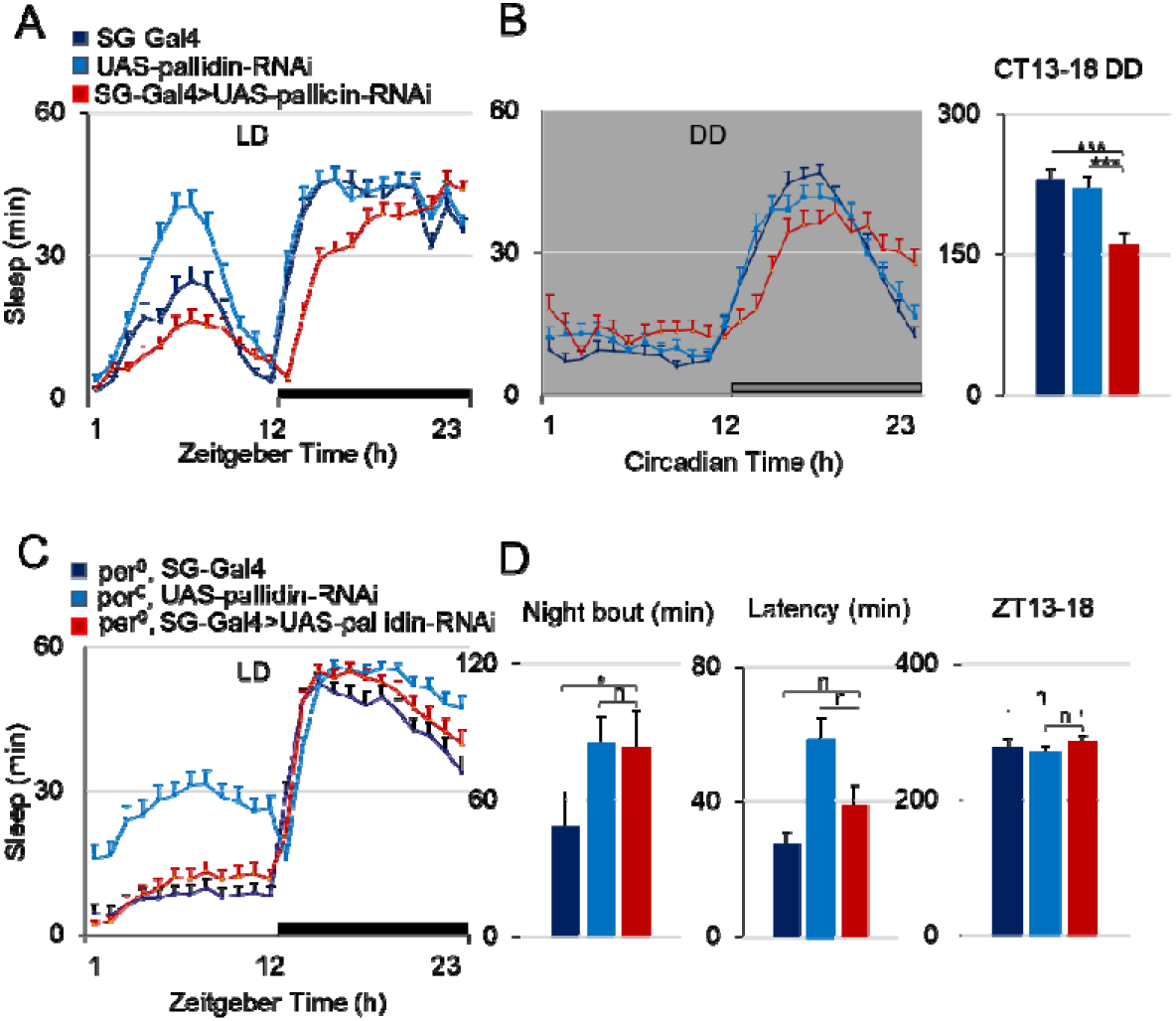
A functional circadian clock is required to observe the sleep/wake phenotype of *pallidin* downregulation in surface glia. (A-B). Baseline daily sleep of *pallidin* knockdown flies in Light Dark condition (LD, A), and following transfer to constant darkness (DD, B). *pallidin* knockdown flies displayed reduced sleep during the presumptive night in constant darkness. C-D) Combining the *period* mutant with *pallidin* downregulation prevents nighttime sleep loss (C), fragmentation or longer latency (D), *: p<0.05, ***: p<0.0005; Kruskal-Wallis with post hoc comparisons between control and knockdown conditions, N=30-43.

### Pallidin is required for sustained GABAergic neuronal activity

In the first half of the night, GABAergic input to the large ventral lateral clock neurons (lLNv) is playing a critical role in promoting sleep^61–64^. To assess the implication of GABAergic transmission in the *pallidin* knockdown sleep phenotype, we fed the flies with the GABA-A agonist 4,5,6,7-tetrahydroisoxazolo-[5,4-c]pyridine-3-ol (THIP), known to induce sleep in *Drosophila*. This treatment potently increased sleep as previously reported^65, 66^, and *pallidin* knockdown flies were no longer significantly different from the controls, suggesting that the phenotype originate from impaired release of GABA (Figure 4A). To further evaluate the implication of GABAergic transmission in the *pallidin* knockdown sleep phenotype, we expressed the heat-dependent cation channel dTrpA1 with the LexA/LexAOP system in GABAergic neurons while downregulating *pallidin* in surface glia. After transfer from 18°C to 25°C the vGAT-lexA>LexAOP-TrpA1 flies displayed reduced latency and larger nighttime sleep amounts as expected following and increase in GABAergic tone (Figure 4B, supplemental Figures S8), while raising the temperature to 30°C induced lethality in those flies. *Pallidin* down-regulation in surface glia completely prevented this effect, suggesting that it is required for the sustained activity of GABAergic neurons.

**Figure 4:**
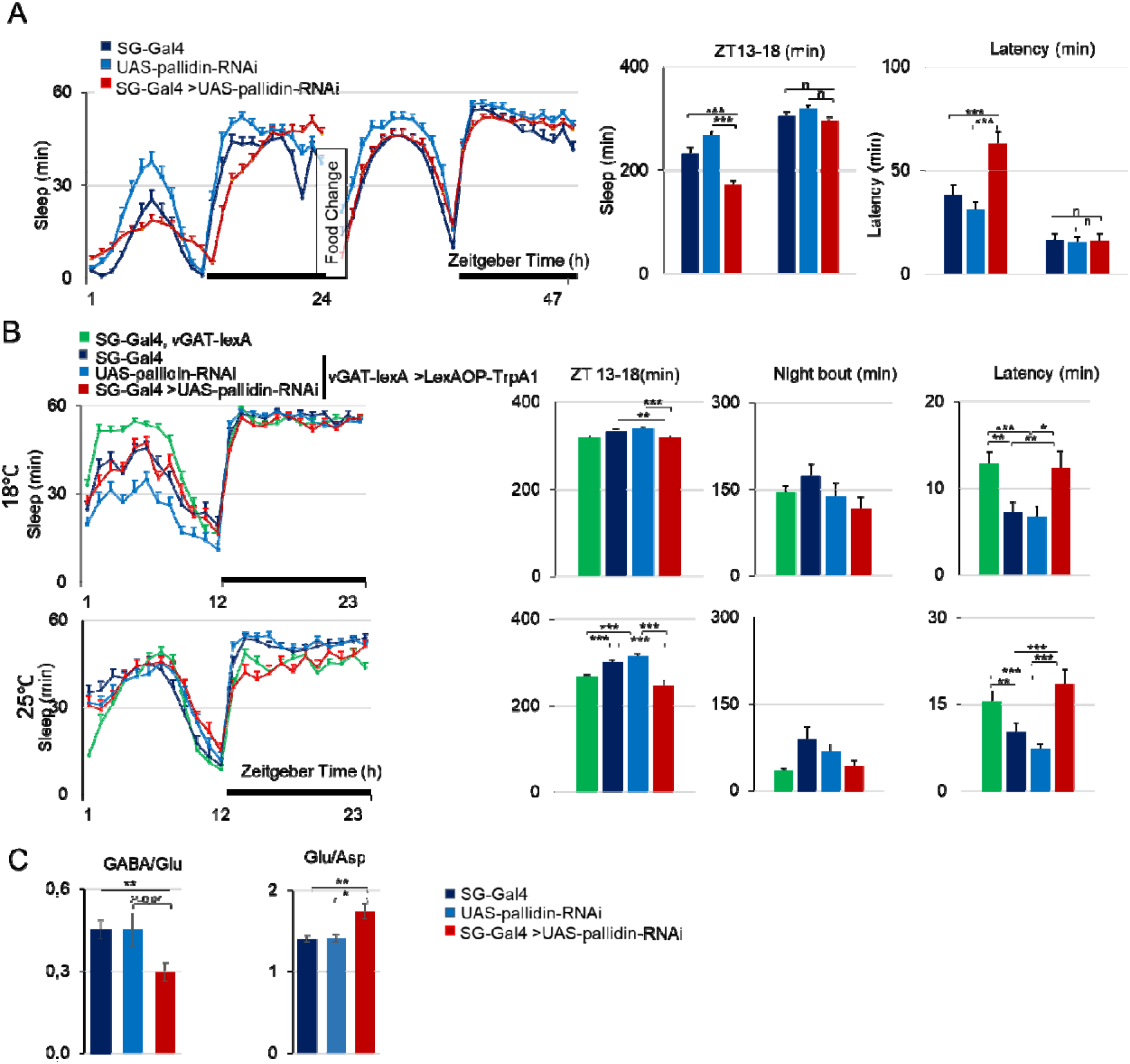
Effect of Pallidin knockdown in surface glia on GABAergic transmission. A) *Pallidin* knockdown flies (in red) and genetic controls (in blue) recorded on normal food, then transferred to food supplemented with 0,1mg/ml Gaboxadol (THIP) for 2 day. The amount of sleep increased in all groups. Hourly sleep, latency, and sleep at ZT13-18 in the knockdown flies become similar to the control groups. ***: p<0.0005; Kruskal-Wallis with post hoc comparisons between control and knockdown conditions. N=45-53. B) Expression of TrpA1 in GABAergic neurons using the vGAT-lexA transgene combined with LexAOP-TrpA1 results in higher sleep amounts and longer sleep bout duration at 25°C, compared to non-TrpA1 expressing controls (green), but not at 18°C (upper panel) when the channels are not activated. *Pallidin* knockdown flies (red) do not display higher night sleep amounts and shorter sleep latency compared to non-TrpA1 expressing flies (green), in contrast to the genetic background controls (dark and light blue). Similar results were obtained with an independent *UAS-pallidin-RNAi* transgene (Supplemental Figure S8). *: p<0.05; **: p<0.005; ***: p<0.0005; Kruskal-Wallis with post hoc comparisons. N=37-43. C) HPLC analysis of GABA, glutamate (Glu), and aspartate (Asp) contents in individual brains. The ratio of GABA to glu concentrations is slightly but not significantly decreased in *pallidin*-downregulation flies(red) compared to the genetic control (blue). The ratio of Glu to Asp was significantly increased in *pallidin*-downregulation . *: p<0.05; **: p<0.005; Kruskal-Wallis with pos hoc comparisons between control and knockdown conditions, N=8-12.

GABA is synthesized by the decarboxylation of glutamate, which is also an excitatory neurotransmitter. Using individual brain HPLC, evaluation of the global content of GABA, glutamate and aspartate, an amino acid metabolically linked to glutamate, showed a trend for a reduction of the GABA/glutamate ratio and an increase in the Glutamate/Aspartate ratio (Figure 4C) indicating that neurotransmitters involved in excitation/inhibition balance are affected by *pallidin* knockdown in surface glia. Surface glia could modulate GABAergic transmission through a diversity of potential mechanisms. Since a previous anatomical description reported subperineurial glia contacts with a subset of neuronal cell bodies^67^, we used the GFP Reconstitution Across Synaptic Partners approach ^68, 69^, to reveal direct cell-cell contacts between surface glia and GABAergic neurons. As shown in Figure 5, we observe extensive cellular contacts between the cell bodies of numerous GABAergic neurons and surface glia.

**Figure 5:**
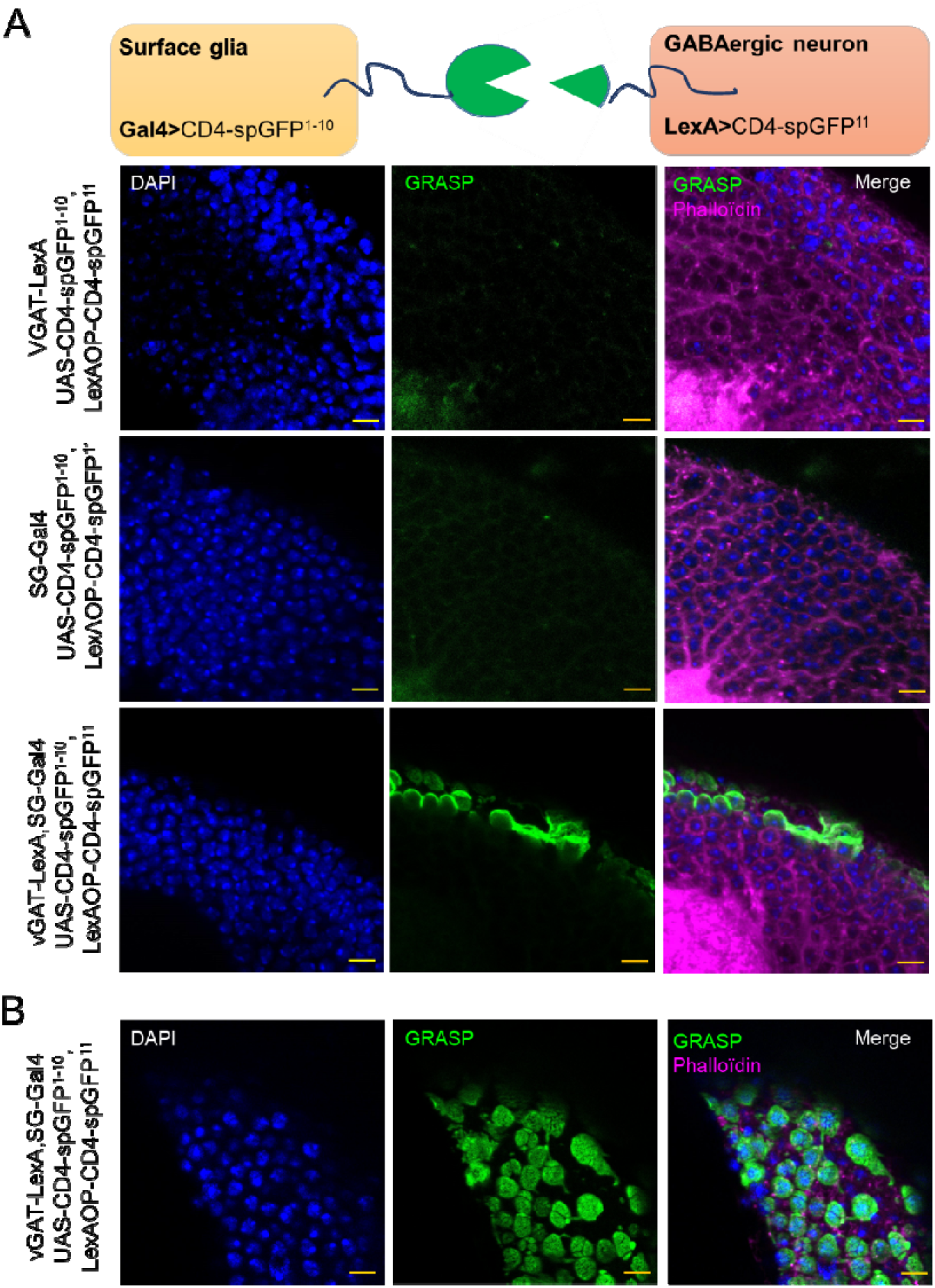
Direct contacts between surface glia and GABAergic neurons. A) Schematic of the GRASP system (see methods) and confocal images (Z stack, maximal intensity projection of 5 images, 1μm apart) of cross sections in the dorsal brain for each genotype tested. GFP fluorescence is observed on the top half of the GABAergic neurons when both surface glia and GABAergic neurons express the GRASP constructs. B) Tangential view of the GRASP signal between GABAergic neurons and surface glia (Z stack, maximal intensity projection of 2 images, 1μm apart). Scale bar: 5μm.

### Other members of the BLOC1 complex genes are required in SG for sleep/wake regulation

To determine whether the sleep-regulatory function of *pallidin* in SG is shared by other BLOC1 genes, we used available UAS-RNAi lines to inhibit their expression in SG. As shown in Figure 5, down-regulation of several BLOC1 members presented a phenotype similar to that seen with *pallidin: dysbindin, Blos1 and Blos2* (Figure 6A-C), while the down-regulation of *snapin* did not affect significantly sleep/wake (Figure 6D).

**Figure 6:**
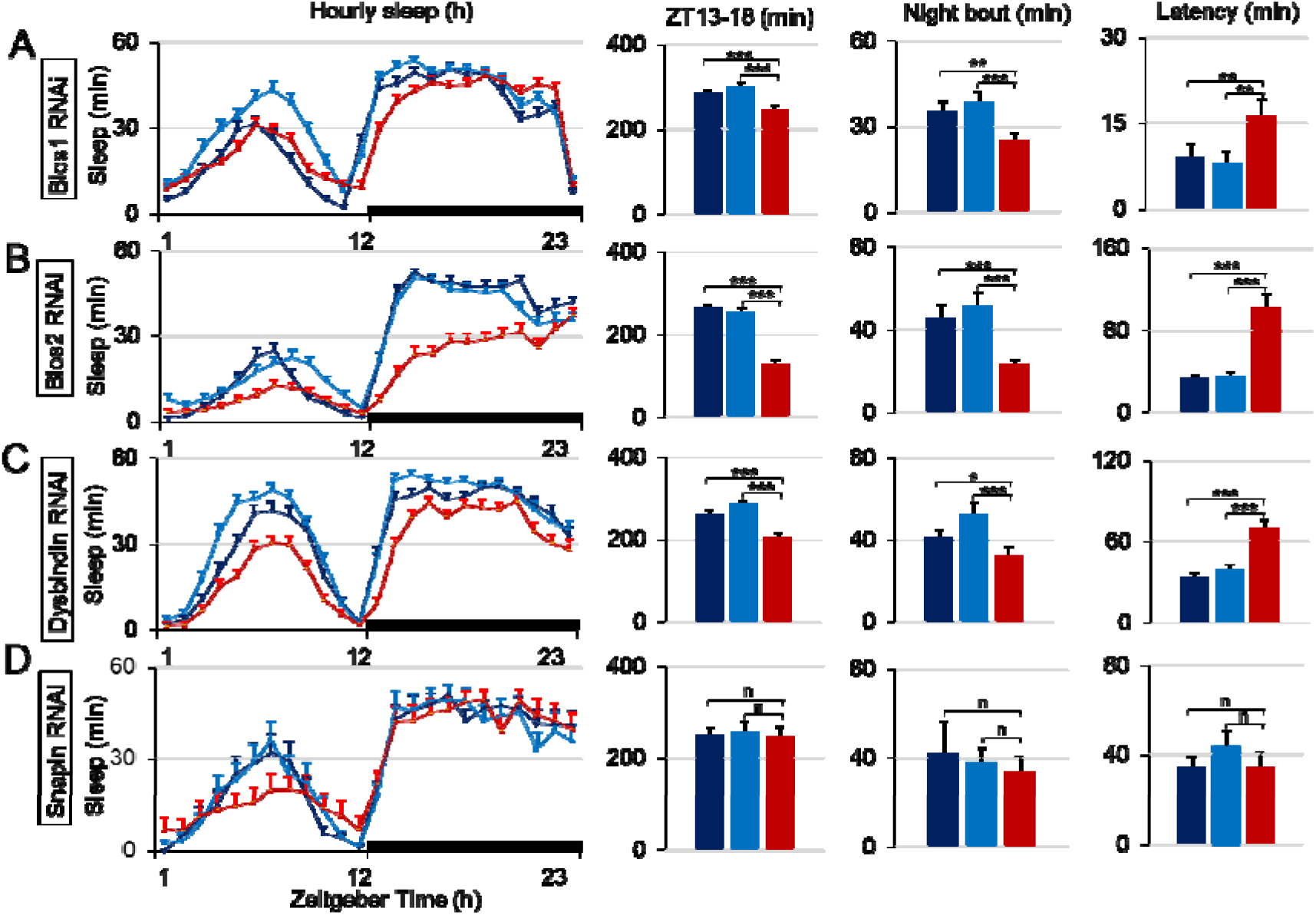
Downregulation of BLOC1 complex genes in surface glia. **A-D**) Baseline daily sleep of flies expressing UAS-Blos1-RNAi, UAS-Blos2-RNAi, UAS-Dysbindin-RNAi, UAS-Snapin-RNAi in surface glial cells (in red) compared to the respective genetic control groups (in blue). In former three cases, nighttime sleep is reduced, delayed (longer latency) and fragmented (shorter average bout duration). *: p<0.05; **: p<0.005; ***: p<0.0005; Kruskal-Wallis with post hoc comparisons between control and knockdown conditions. N=42-52, except N=13-15 for snapin.

A mutation in the pallidin gene, *pallidin*^Δ^*^1^*, devoid expression in the larva, has been recently described^9^. *pallidin*^Δ^*^1^* homozygous flies or in which *pallidin^Δ^*^1^ is combined with a deficiency covering the *pallidin locus*, displayed longer sleep time throughout day and night, and shorter latency (Supplemental Figure S9). These results are difficult to compare with the mild and selective knockdown provided by the Gal4/UAS system, but nonetheless provide further evidence that pallidin is involved in sleep/wake regulation.

### LAT1-like transporters and TOR signaling knockdown in surface glia affects early night sleep

The function of pallidin in the mouse brain has been linked to the transport and/or the availability of large neutral amino acids^34, 35^, which are controlled to a large extend by the LAT1 transporter localized in blood-brain-barrier cells^70–74^. We therefore conducted a set of experiments in *Drosophila* to test the hypothesis that the sleep phenotype observed upon *pallidin* down-regulation could be linked to amino acid availability in the brain. We first found that inhibition of the *Drosophila* homologs of LAT1: *JhI-21 (Juvenile Inducible 21), mnd (Minidisc)* as well as *CD98Hc*, the *Drosophila* homolog of the gene coding for the heavy-chain subunit necessary for the expression of a functional transporter, replicates the *SG>UAS-pallidin-RNAi* sleep phenotype (Figure 7A-B, Supplemental Figure S10). Secondly, inhibition of the TOR (Target of Rapamycin) cellular amino acid sensing pathway using *UAS-TOR-RNAi* or a *UAS-raptor-RNAi* resulted in the same effect (Figure 7C-D). These results suggest that LAT1-like transporters, TOR and Pallidin could participate in a process regulating both amino acids intake and sleep. In support of this idea, the intake of Leucine through LAT1 is required to activate TOR signaling in some cellular contexts^75^.

**Figure 7:**
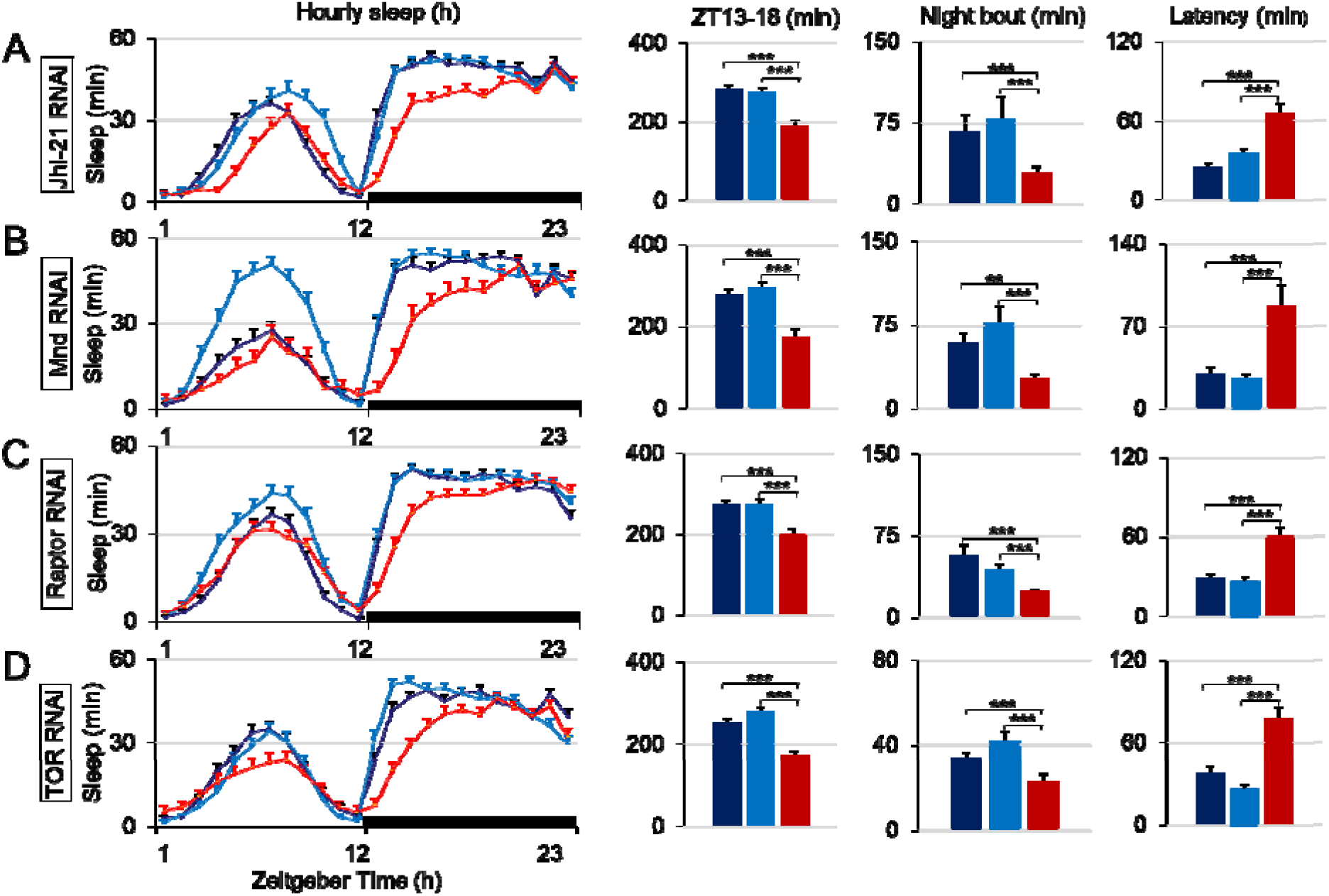
Downregulation of LAT1-like homologues and TOR, Raptor in surface glia. Baseline daily sleep of flies expressing UAS-JhI-21-RNAi, UAS-Mnd-RNAi, **(A-B)** UAS-Raptor-RNAi, UAS-TOR-RNAi (**C-D**) in surface glial cells (in red) compared to the respective genetic control groups (in blue) In all cases, latency was increased and night sleep reduced compared to controls. Average night bout was also reduced except for the TOR-RNAi expressing flies. *: p<0.05; **: p<0.005; ***: p<0.0005; Kruskal-Wallis with post hoc comparisons between control and knockdown conditions, N=44-46.

### Leucine supplementation normalizes the pallidin and LAT1-like transporters knockdown phenotypes

To test the hypothesis that pallidin knockdown flies suffer from a deficit in amino acids, notably essential amino acids that are provided by feeding, we attempted to rescue the phenotype with a nutritional approach. We find that supplementing food with Leucine rescues the *SG>UAS-pallidin-RNAi*, the *SG>UAS-JhI-21-RNAi* and *SG>UAS-Blos2-RNAi* sleep phenotypes (Figure 8, Supplemental Figure S11), while having little or no detectable effect in the genetic controls. The effect of amino acid supplementation was prominent on night sleep, reducing latency, increasing sleep amounts and sleep bout duration. These results link the *SG>UAS-pallidin-RNAi* sleep phenotype to amino acid uptake. Supplementing food with other essential amino acid such as Valine or Tryptophan rescued the *JhI-21* knockdown but, interestingly, not pallidin knockdown (Supplemental Figure S12). As mated females flies have recently been shown to eat more amino acids at night, a behavior absent in males and in non-mated females^76^ we evaluated the *SG>UAS-pallidin-RNAi* night sleep phenotype in mated females and in males. The sleep phenotype was unaffected by the mating status of females and was observed in males, with the exception of longer latency for the latter (supplemental Figure S13).

**Figure 8:**
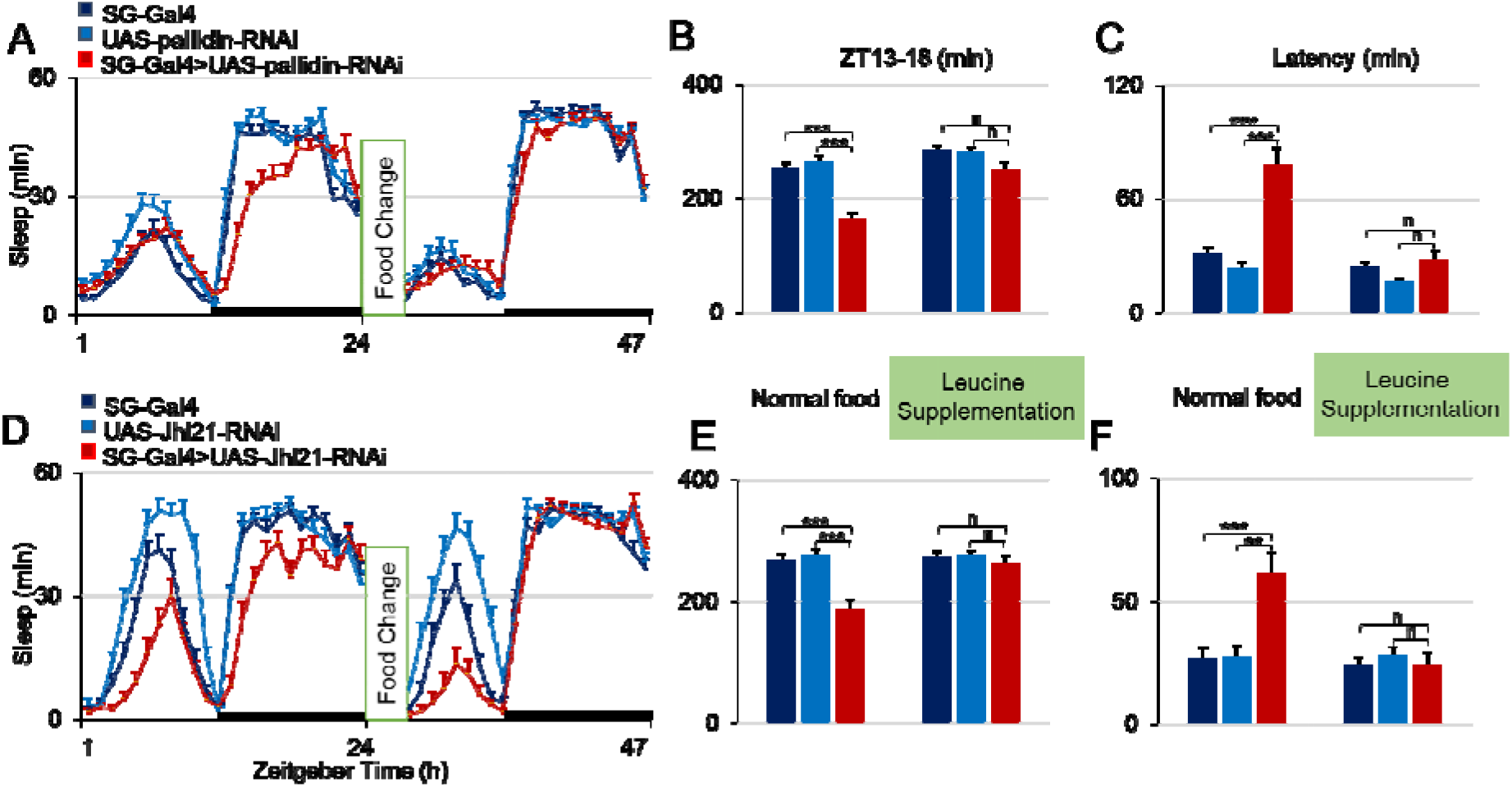
Food supplemented with leucine normalizes the sleep phenotypes of *Pallidin* and *JhI-21* downregulation in surface glia. **A-C**) Pallidin knockdown flies (in red) and genetic controls (in blue) recorded on normal food, then transferred to normal food or normal food supplemented with 50 mM leucine for 1 day. Hourly sleep, latency, and sleep at ZT13-18 in the knockdown flies become similar to the control groups. **D-E**) The same protocol was applied to *JhI-21* knockdown flies and similarly normalized nighttime sleep. *: p<0.05; **: p<0.005; ***: p<0.0005; Kruskal-Wallis with post hoc comparisons between control and knockdown conditions, N=29-45.

### Pallidin down-regulation result in an abnormal trafficking of the LAT1-like *JhI-21* transporter

Since pallidin and the BLOC1 complex can regulate transmembrane protein trafficking and in particular transporters^77, 78^, we evaluated the subcellular localization of *JhI-21* in surface glia using a validated antibody^79, 80^. To unambiguously identify perineurial and subperineurial glia cells, we labelled their nuclei by the expression of a nuclear-βgalactosidase (Figure 9A) or by the expression of a *UAS-GFP-mcherry-ATG8* transgene to label both the nuclei and lysosomal organelles, including autophagosomes^81^ (Figure 9B). *JhI-21* was detected in punctae in the perinuclear region of perineurial and subperineurial cells, as well as in neurons, and colocalized partially with GFP-mcherry-ATG8 suggesting that at least a subset of the transporter is internalized in lysosomes. We observed that the knockdown of *pallidin* in surface glia resulted in an accumulation of *JhI-21* immunofluorescence in punctae within the nuclei of both perineurial and subperineurial glia, suggesting an abnormal trafficking of the transporter (Figure 9A-D). *Pallidin* knockdown did not result in major changes in autophagy as detected with the *UAS-GFP-mcherry-ATG8* construct^81^ (Supplemental Figure S14).

**Figure 9:**
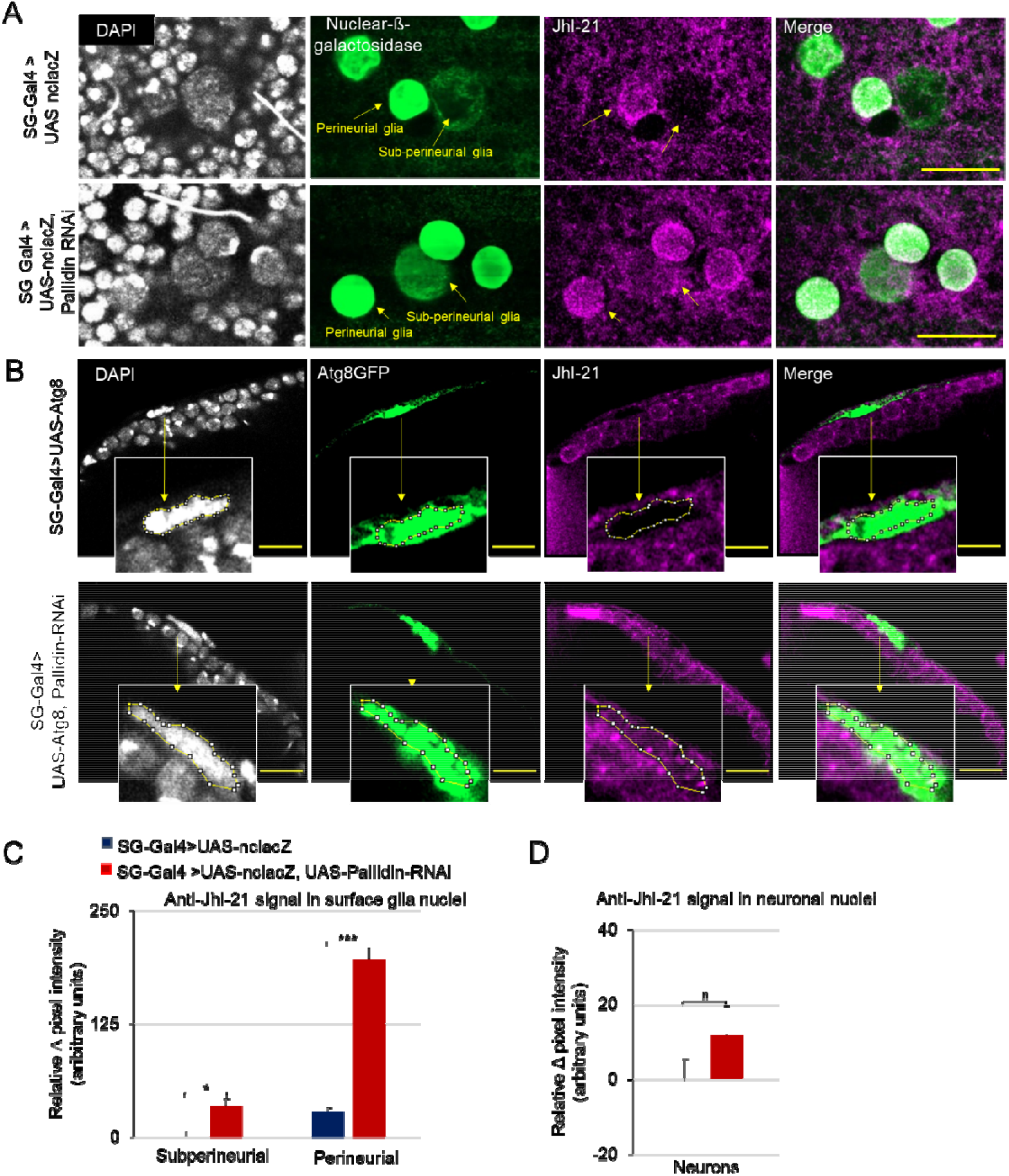
The knockdown of *pallidin* modifies JhI-21 subcellular localization in surface glia. **A**) Tangential sections of the surface of brains expressing either the *UAS-nuclear-LacZ (UAS-ncLacZ)* alone or together with *UAS-pallidin-RNAi* under the control of *9-137-gal4*. Perineurial and subperineurial cells are identified by virtue of the size and location of the anti-ß-galactosidase labelled nuclei (green). The anti-JhI-21 antibody reveals a substantial signal within the nuclei of both perineurial and subperineurial glia of *UAS-pallidin-RNAi* expressing brains, not observed in the control condition. **B**) Cross-section images showing a single subperineurial cell nucleus in brains expressing either the *UAS-GFP-mcherry-ATG8* construct alone or together with *UAS-pallidin-RNAi* under the control of *9-137-Gal4*. GFP-mcherry-ATG8 expression is revealed by an anti-GFP antibody and is present both in the cytoplasm and in the nucleus of surface glial cells. The anti-JhI-21 antibody signal is absent from the nucleus in the control condition but detected as punctae double labelled with ATG8-mcherry-GFP within the *UAS-pallidin-RNAi* expressing flies. (A-C) Bar =10μm, N=4-8. Single confocal sections are shown. **C**) Quantification of the anti-JhI-21 signal in the nuclei of subperineurial (N=8-10) and perineurial (N=28-36) cells of the brains represented in (B). **D**) Quantification of the anti-JhI-21 signal in the nuclei of neurons (N=C18-24) shows no significant difference between pallidin knockdown and control conditions. *: p<0.05; **: p<0.005; ***: p<0.0005; Mann-Whitney between control and knockdown conditions.

### Pallidin in surface glia does not modulate dopamine synthesis

Although the data outlined so far suggest a role for *pallidin* in the regulation of GABAergic transmission, previous reports suggested that this gene could modulate monoaminergic transmission. In particular, it has been shown previously that mice with a defective *pallidin* gene were less sensitive to intraperitoneal injections of tryptophan and of L-DOPA, as evidenced by the reduced behavioral effects and the lesser increase in brain serotonin and dopamine levels, respectively, suggesting that the gene could be required to provide the brain with adequate levels of precursors for monoamine synthesis^35^. A similar hypothesis has been proposed for dysbindin, which can regulate *Drosophila* brain serotonin and dopamine levels, although in the latter case, a deficit in dopamine recycling has been proposed ^27, 30, 82, 83^. To test whether dopamine synthesis could explain the sleep/wake phenotype of *SG>UAS-pallidin-RNAi* flies we conducted similar experiments in Drosophila. Feeding flies with L-DOPA leads to nocturnal insomnia within 24h and a large increase in dopamine synthesis^80^. The sensitivity to L-DOPA as evaluated by sleep loss was not significantly different between *SG>UAS-pallidin-RNAi* flies and controls, while food intake remained equal in all the groups (Supplemental figure S15-A). Furthermore, the global levels of dopamine measured by HPLC were increased to a similar extend in *pallidin* knockdown flies compared to the control groups (Supplemental figure S15-B). We observed no difference in dopamine or 5-HT levels in *SG>UAS-pallidin-RNAi* in the vehicle control condition. To further test the involvement of surface glia and *pallidin* in dopaminergic transmission, we combined the knockdown of *pallidin* with the *fumin* mutation of the *dopamine transporter (DAT)* gene. This mutation impairs the reuptake of dopamine, and leads to a hyperdopaminergic state and severe insomnia^84, 85^. Given the lack of dopamine reuptake, it can be assumed that dopaminergic transmission in this mutant relies heavily on de novo dopamine synthesis^86^. It could then be assumed that *pallidin* knockdown would attenuate the *fumin* phenotype by preventing the transport of amino acids. *Pallidin* down-regulation in surface glia failed to attenuate the severity of fumin-driven insomnia, suggesting that it does not impact dopaminergic synaptic transmission directly (Supplemental figure S15-C).

## Discussion

We provide evidence in drosophila that Pallidin and other components of the BLOC1 complex regulate the initiation and maintenance of sleep in the early part of the night, at the level of surface glia cells, the equivalent of the BBB. The data support a mechanism whereby BLOC1 regulates LAT1-like transporters subcellular trafficking in both perineurial and subperineurial cells, leading to adequate essential amino acid supply and GABAergic neuronal activity promoting sleep, in a circadian clock dependent manner.

Sleep during the night is qualitatively different from the light phase with higher arousal thresholds and longer sleep bout duration^46, 59^. This is particularly relevant for the first half of the night, where flies experience the longest bout of sleep of the day^45^. These differences presumably result from the homeostatic sleep drive accumulated during the light phase and from circadian factors favoring sleep during the early night. The delayed, reduced and fragmented night sleep observed upon *pallidin* knockdown in surface glia may thus originate from a failure to sense homeostatic sleep pressure, a failure to elicit deep sleep, or from circadian clock related deficits. The assessment of the sleep homeostatic response to sleep deprivation and of sleep intensity (Supplemental Figure S6-7) suggests that the down-regulation of *pallidin* does not prevent flies from experiencing deep sleep or compensating for sleep loss, making these processes unlikely to explain by themselves the observed sleep phenotype. Similarly, *pallidin* knockdown flies experienced normal sleep/wake circadian rhythms and display near 24h circadian period under constant dark conditions (Figure 2), demonstrating that the core circadian time-keeping system is functional. Conversely, blocking the cycling of the molecular clock by placing flies in a per^0^ mutant background completely abolishes the *pallidin* knockdown sleep/wake phenotype, showing that the circadian clock is implicated in this *pallidin*-dependent process. This raises the possibility that *pallidin* affects one or several clock dependent sleep/wake regulatory networks, and in particular those under the control of the large PDF expressing neurons (lLNv). In these neurons, GABAergic input has been shown to be modulated in a circadian manner resulting in higher lLNv inhibition in the early night, promoting sleep, while a disruption of this input results in delayed and fragmented sleep^62–64, 87^. This would explain the tight coupling we observe between sleep loss in the early part of the night and increased latency to nighttime sleep. Since the longest sleep bouts of the 24h day occur usually in the first part of the night, this also explains the reduced average sleep bout duration in the knockdown flies. The GABAergic neurons responsible for this inhibition are so far unidentified. The delayed and reduced night sleep observed upon *pallidin* knockdown in surface glia is also similar to the phenotype previously reported with pan-GABAergic neuron inhibition^88^. Conversely, feeding flies the GABA agonist THIP^65^ or increasing GABAergic neuronal firing by expressing TrpA1 (Figures 4) can potently induce sleep. We find here that the latter manipulation cannot promote sleep when *pallidin* is downregulated in surface glia, suggesting that the gene is required to allow sustained GABAergic neuronal activity. Alternatively *pallidin* function could be regulated directly by the clock present in perineurial cells, which modulates the efflux properties of subperineurial cells in a circadian manner^89^.

Down-regulation of several members of the BLOC1 complex: Blos1, Blos2, and dysbindin, in surface glia phenocopy the knockdown of *pallidin*, strongly suggesting that the BLOC1 complex is involved in this *pallidin*-dependent sleep/wake regulation. This is consistent with the observation that the pool of *pallidin* and dysbindin present in the cell is almost entirely associated with the complex^1, 36^, and that the lack of one subunit leads to a destabilization of the others (for example ^9, 10^). The lack of sleep alteration with snapin manipulation may result from lower efficiency of the genetic tools used for down-regulation. Alternatively, it may also reflect different gene dosage sensitivity and functionality for the different members of the complex, as previously reported^8, 9, 29^, or the existence of multiple subunits within the complex^2^, or different complexes containing a subset of the subunits^20^. In *Drosophila* larval neuromuscular junction, Snapin has for example been shown to regulate synaptic homeostasis^26^, while Pallidin does not seem to affect baseline neurotransmission, and is required during sustained neuronal activity to replenish the pool of releasable synaptic vesicles ^9^. The BLOC1 complex has been shown to be required for the biogenesis of recycling endosomes, through its interactions with sorting endosomes and the cytoskeleton^4^. These BLOC1 functions and the regulation of sleep by BLOC1 members reported in this study are in agreement with a recent report showing bidirectional interactions between sleep and endocytosis in surface glia^52^. Intriguingly, the endocytosis in surface glia was reported to be the most intense in the early part of the night^52^, where sleep is the deepest and the phenotype of *pallidin* down-regulation is the most pronounced. In addition, the activity of the recycling endosome associated small GTPase rab11 appears to play a prominent role in this context^52^. Consistent with these findings, extensive protein-protein interactions analyses identified rab11 as the best interacting partner for BLOC1 in both *Drosophila* and humans^5^. Thus, BLOC1 may facilitate the high endocytic activity of surface glia during the early part of the night, notably the biogenesis of recycling endosomes, while having a less prominent role at other times during the day.

Flies completely defective for the *pallidin* gene did not display the same phenotype as the one obtained with the surface glia specific knockdown. As *pallidin* is expressed at all developmental stages and in most if not all cell types, including neurons and glial cells within the brain^90–92^, the mutation is likely to have a compound effect, making the interpretation of the adult sleep/wake phenotype difficult to interpret. In contrast and in line with our results, *pallidin* mutant mice display reduced sleep amount during the light phase of the day, the primary sleep period in rodents, and shorter average sleep bout duration^32^. However, BBB specific and conditional knockout models in mice would be necessary to determine whether BLOC1 is similarly regulating sleep/wake in mammals. The incidence of sleep-wake disorders have not been reported for humans suffering from the rare Hermansky–Pudlak syndromes, bearing deficits in BLOC1 or other functionally related complexes, but would deserve scrutiny^93^. We identify here a new role for this complex in the interface between the brain and the circulatory system, affecting sleep and potentially cognitive processes, including schizophrenia and psychiatric disorders that may be linked to abnormal sleep/wake regulation. The role of the mammalian BBB in sleep/wake regulation is virtually unexplored, however whole brain calcium imaging during sleep in the zebrafish larvae indicate that ependymal cells could be a significant component of the sleep regulatory system^94^.

Surface glial cells express tight junctions, and molecular components in common with the mammalian BBB including claudins and homologous efflux transporters. As in the BBB this organization prevents the entry of macromolecules and tightly regulates the exchanges of solutes between the circulatory system (the hemolymph in drosophila) and the brain, thus maintaining the particular interstitial fluid composition necessary for neuronal activity^53^. This function relies on intense transporter activity. Early studies of *pallidin* function pointed to a role in amino acid import into the brain transport as suggested by lower sensitivity to intraperitoneal injections of the LAT1 substrates L-DOPA and tryptophan^35^. The LAT1 transporter plays a major role in the import of these amino acids, and in the import of large neutral essential amino acids such as leucine, isoleucine and histidine^95^. It is prominently expressed in the endothelial cells of the brain capillaries ^72, 96^. A conditional knock out of LAT1 in brain endothelial cells confirmed the crucial role of LAT1 in the regulation essential amino acid abundance within the brain, its impact on GABAergic transmission and its potential implication in autism spectrum disorders^97^. Intriguingly, the relative abundance of amino acids in the brain of *pallidin* mutant mice resembles those found in mice with a BBB specific knock-out LAT1^34^. Here, we provide evidence that *pallidin* function in surface glia is linked to essential amino acid supply and to LAT1-like transporter activity. First, down-regulation of the LAT1-like transporters *JhI-21* and minidisc, the heavy subunit CD98Hc and the TOR signaling phenocopy completely or to a large extend the BLOC1 genes knockdown. Second, supplementing the food with leucine can normalize the JhI-21, *pallidin* and Blos2 phenotypes, further implicating amino acid supply. Interestingly, the phenotype of the *JhI-21* transporter knockdown were rescued by valine and tryptophan supplementation, while *pallidin* knockdown could only be rescued by leucine. This suggests that Pallidin does not solely modulate LAT1-like transporter function, and has complex effect on multiple transport systems, as previously suggested ^34, 35^. *JhI-21* and *minidisc* are the *Drosophila* closest homologs of LAT1^77^. *JhI-21* is expressed broadly in the *Drosophila* brain, with most of the signal in perinuclear punctae, that colocalize partially with the lysosomal marker ATG8 (Figure 9). The subcellular trafficking of *JhI-21* in surface glia is abnormal following *pallidin* down-regulation, with a substantial fraction of the transporter localized in the nucleus (Figure 9). This abnormal trafficking is likely to reduce the functionality of the transporter, explaining the similar sleep/wake phenotypes of *JhI-21* and *pallidin* downregulation in surface glia and their normalization by essential amino acid supplementation. A nuclear localization has already been reported in glioma cell lines for the Solute Carrier (SLC) Eaat1 and Eaat2 transporters^99,100^, and for the K+ Inwardly rectifying channel^99^. In both cases, this unusual subcellular localization was associated with an overall reduced functionality of the protein in the cell. Aside transporters, several full length transmembrane receptors have been repeatedly reported to be localized in the nucleus where some of them may act as a transcription factors^101,102^. The mechanisms underlying these phenomenon are mostly unknown. In this study, we were not able to detect obvious changes in autophagic flux, suggesting that *pallidin* down-regulation does not induce major changes in lysosomal activity, and rather affects more specifically particular cargos. In mammalian epidermal cells BLOC1 function does not affect the trafficking of all protein and rather results in the accumulation of particular sets of cargo in specific subcellular compartments, notably tyrosinase-related protein-1 and the ATP7 copper transporter^4, 12, 78^. How such an accumulation would lead to nuclear localization in surface glia remains to be determined. Interestingly, in addition to their cytoplasmic localization, a nuclear localization has previously been reported for two BLOC1 subunits: BLOS2^103^ and in particular dysbindin^82^. The latter was for instance found in the nuclei of all neurons in postmortem human brains^21^, and in cell cultures where a report suggested that it could shuttle in and out to regulate the expression of synapsin^104^. Interestingly in this context, recent report has also shown that ATG8 is present both in lysosomes and in the nucleus where it can regulate gene expression in association with transcription factors^81, 105^. Such lysosomal-nuclear connections open the possibility of trapping a diversity of unexpected proteins in the nucleus in normal as well as in abnormal conditions.

The data presented here emphasize the implication of circulating amino acids and in particular essential amino acids in sleep/wake regulation, corroborating several recent metabolomics studies in mammalian models and in human^106, 107^ where branched chain amino acids (BCAA) levels have been repeatedly shown to be modulated by the circadian clock and/or the sleep homeostat. For example, a recent study identified in the plasma of insomniac patients prominent changes in the levels of several BCAA, including increased levels of leucine during the night^108^. Food supplemented with BCAA can correct sleep disorders in a mouse model with chronic sleep disruption^109^. Amino acids are at the interplay between many processes, being involved in protein synthesis, energy metabolism, neurotransmitter synthesis and degradation. Amino acid transporters may further intertwine these processes: as a prime example, LAT1 is an antiporter that can associate the import of leucine to the export of glutamine, a crucial metabolite for the recycling of glutamate and GABA in the brain^75^. Interestingly, we find here that the GABA/ glutamate ratio tend to be decreased in *pallidin* knockdown flies while the glutamate/aspartate ratio is increased, compared to controls. These results suggest that BLOC1 function in surface glia modulates neurotransmitter and brain amino acid metabolism on a global level, and/or to control the inhibitory/excitatory balance. In Drosophila, protein intake, threonine intake and D-serine levels play a role in sleep/wake regulation^55, 110–112^. While the changes in amino acid intake presumably change global free amino acid levels within the brain, the effect on sleep/wake regulation seem to originate from specific sleep/wake regulatory networks or neurotransmission systems. D-serine appear to act via NMDA receptor transmission to upregulate sleep^110^, and the modulation of sleep by protein intake implicate ilp2 producing neurons^55^. Threonine supplementation promotes sleep in a clock independent manner, by inhibiting metabotropic GABA receptors on R2 sleep promoting neurons^111^. In humans, amino acids supplemented diets, notably with tryptophan or tyrosine, have been designed to improve sleep and wakefulness, based on the principle that it would increase the synthesis of serotonin and dopamine, respectively^70, 113–115^. Our results point to a model independent from these previously identified mechanisms: higher recycling endosome activity in the early part of the night, mediated by BLOC1, would lead to high LAT1-like activity and essential amino acid import in the brain. The TOR signaling appear also to be required and may facilitate LAT1-like transporter function. Our assessment of brain dopamine and serotonin levels, and our *fumin*/ L-DOPA data suggest that this *pallidin*-dependent sleep regulation does not involve a modulation of monoamine levels, in contrast with previous reports ^27, 35^. However, in these reports the conclusions were reached using complete knock out and not by down-regulation in specific cell types, making the comparison with the present results difficult. For example, evidence in *Drosophila* suggest that dysbindin regulates global brain dopamine level by modulating its recycling through glial cells^27^. In contrast, our data suggest that essential amino acid import to the brain facilitated by *pallidin* would enhance GABAergic transmission required for sleep. The neuronal networks and the mechanisms involved remain to be elucidated. Incorporation experiments using isotopically labelled leucine have suggested an association between GABAergic neuronal activity and protein synthesis^116^. The converse association was observed in similar experiments assessing the dopaminergic and serotoninergic systems^116^. The GABAergic sleep promoting neurons modulated by *pallidin* may be able to sense essential amino acid such as leucine, as shown for the Ilp2 producing neurons^79^, where leucine uptake generates a calcium signal under the control of *JhI-21* and mnd. During larval development, dietary amino acid have also been shown to trigger calcium waves in subperineurial glia and the release of insulin-like peptides^117^. We show here that a large number of GABAergic neurons in the adult fly brain are in direct cellular contacts with surface glia and could thus receive such peptidergic signals. It has been shown recently that insulin-like peptide released from cortex glia could activate specific energy metabolic pathways in neurons and promote long term memory^118^. Similar mechanisms could be involved in sleep wake regulation. Another hypothesis, not incompatible with those previously mentioned can be put forward: the capacity of LAT1-like transporters to regulate glutamine could increase the flux of GABA-glutamate-glutamine cycle and regulate GABA homeostasis in the interstitial fluid, as suggested from pharmacological experiments^95, 119^. Previous studies in dysbindin mutant mice have shown that BLOC1 can disrupt GABAergic activity^31^ which is also affected in schizophrenia^20^. Interestingly, one of the prominent sleep abnormality in schizophrenia patients is a reduction in sleep spindle density^40^, which strongly relies on GABAergic activity^120, 121^. A potential role for amino acid transport at the blood-brain-barrier in those contexts would deserve investigation given our results. It is worth noting that a conditional LAT1 knockout at the blood-brain-barrier induces autism-like phenotypes that can be normalized by amino acids^97^, and that genes involved in amino-acid transport are among those commonly disrupted in autism and schizophrenia^122^. In conclusion, this study provides potential mechanisms at the blood brain interface that may be relevant to both sleep disruption and psychosis, and emphasize the possibility of diet-based therapies.

## Conflict of interest

The authors declare no conflict of interest

## Acknowledgments

We thank the Bloomington Stock Center, the Vienna *Drosophila* RNAi Center, S. Birman, M. Bourouis, B. Mollereau, and I. Alliaga for fly stocks, Miss Lorie Meunier and Mr. Paul Sauvageot for the HPLC protocol implementation for amino acid content analysis, Serge Birman for the critical reading of the manuscript. This work was funded by INSERM (https://www.inserm.fr), CNRS (https://www.cnrs.fr/fr/page-daccueil), Université Claude Bernard Lyon 1 (https://www.univ-lyon1.fr), région Auvergne-Rhône-Alpes and SFRMS (www.sfrms-sommeil.org) .HL received a scholarship from the China Scholarship Council (CSC, https://www.chinesescholarshipcouncil.com/) and SA received a scholarship from Université de Tunis El Manar (http://www.utm.rnu.tn).

---

**Supplemental figure S1:**
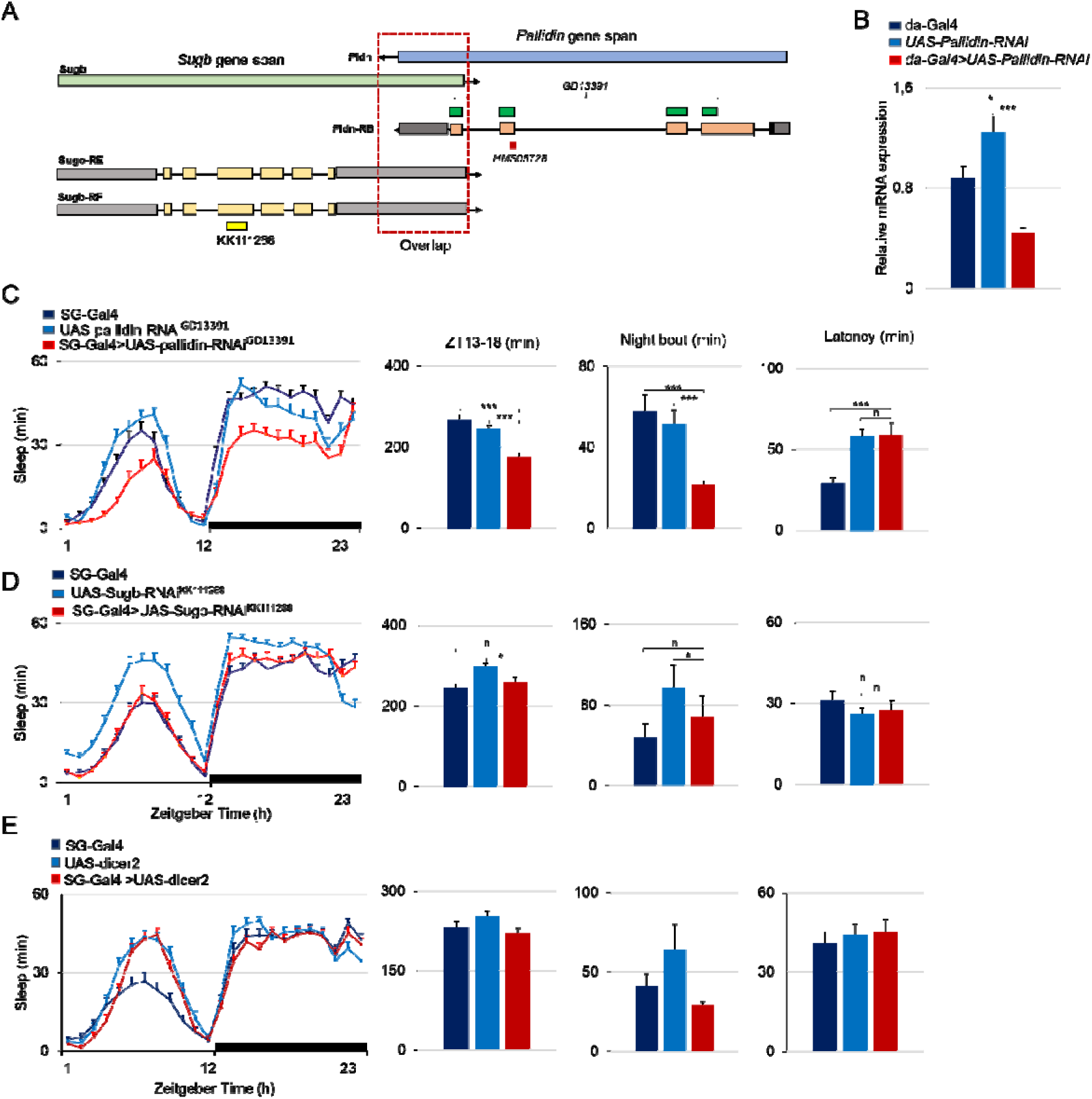
Pallidin UAS-RNAi transgenes, knockdown of *Sugar* baby and over-expression of dicer2. A) Graphical representation of the *pallidin* gene and localization of the areas targeted by the GD13391 (green) and HMS05728 *pallidin-RNAi* constructs. The Pallidin gene shares exon sequences (red dotted square) with Sugar baby included in GD13391(adapted from Flybase, not to scale). B) Relative *pallidin* mRNA levels in the bodies of *da-Gal4>UAS-Pallidin-RNAi*^MS05728^ flies as evaluated by QPCR (values shown: 1000 x (*pallidin/RP49*). * p<0.05, ***, p<0.0005; N=9 (da-Gal4), 9 (*UAS-pallidin-RNAi*), 7 (*da-Gal4>UAS-Pallidin-RNAi*). Kruskal-Wallis with post hoc comparisons between control and mutant conditions. C) *Pallidin* knockdown in surface glia using *UAS-pallidin-RNAi*^GD13391^ (corresponding to the GD13391 construct ID). N=41-47. D) Sugar baby knockdown in surface glia using *UAS-Sugb-RNAi*^iKK111268^: baseline is not affected when compared to controls. N=46-51. E) UAS-dicer-2 expression in surface glia does not disturb sleep. Hourly sleep upon expression of UAS-dicer-2 driven in surface glial cells (ElavGal80,9-137-Gal4 driver) in red, compared to the respective genetic control groups N=45-50. *: p<0.05; ***: p<0.0005; Kruskal-Wallis with post hoc comparisons between control and knockdown conditions.

**Supplemental figure S2:**
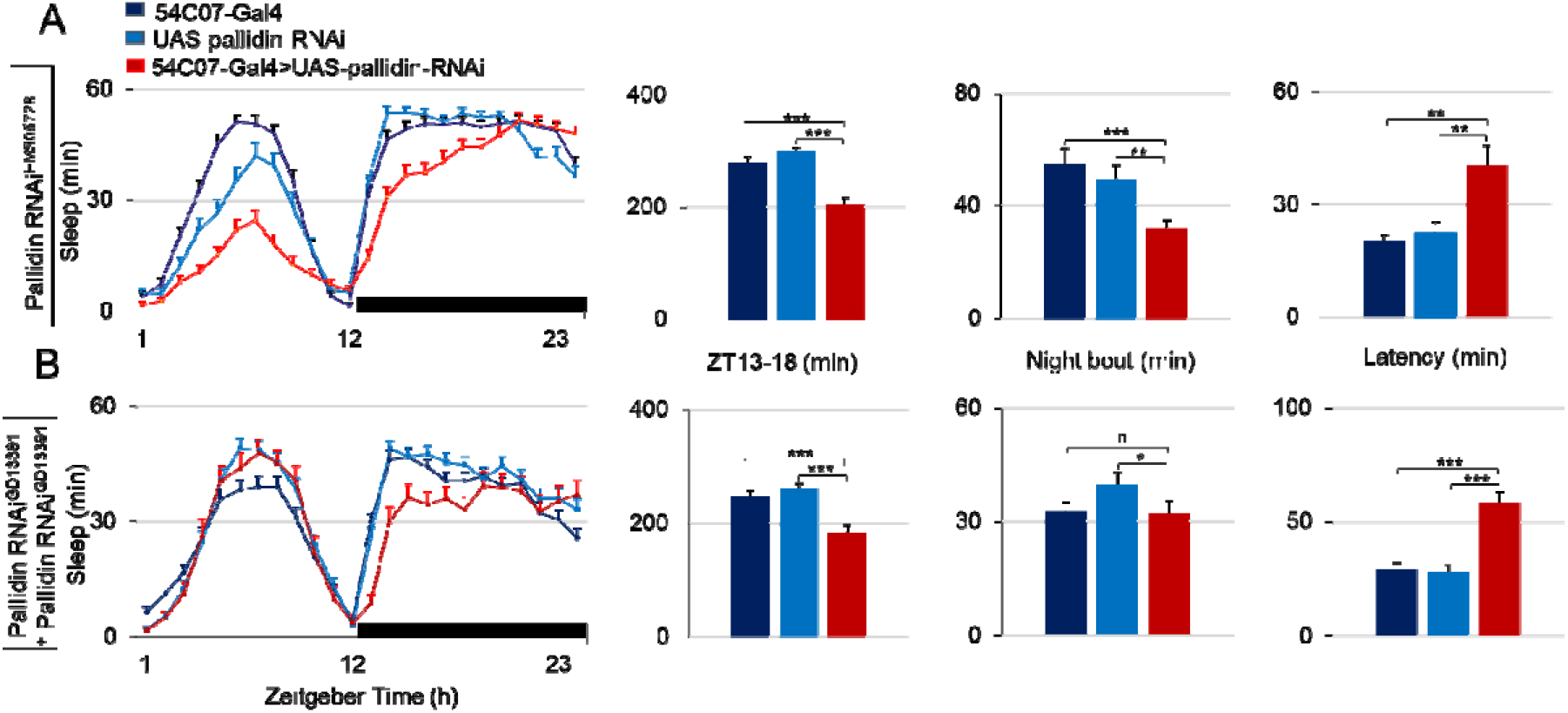
pallidin knockdown in subperineurial-glia reduces nighttime sleep. *pallidin* knockdown in subperineurial glia using the *54C07-Gal4* driver and *UAS-pallidin-RNAi*^HMS05728^ flies (**A**, n=38-48) or *UAS-pallidin-RNAi*^GD23322-3^ (**B**, n=48-55) causes reduced sleep time, especially in the first half night (ZT13-18) and long latency. *: p<0.05; **: p<0.005; ***: p<0.0005; Kruskal-Wallis with post hoc comparisons between control and mutant conditions.

**Supplemental figure S3:**
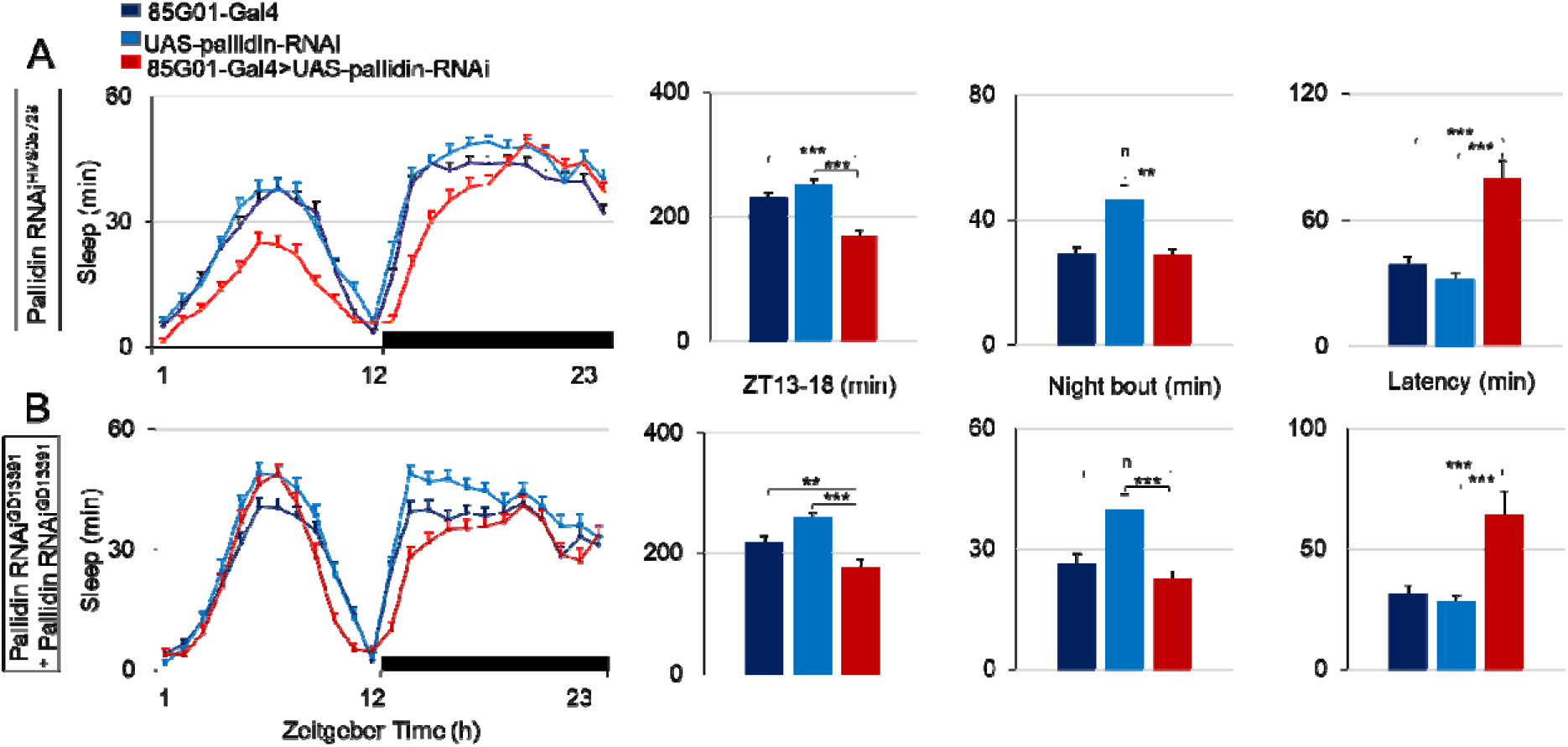
*pallidin* knockdown in perineurial-glial reduces nighttime sleep. Knockdown of *pallidin* using the *85G01-Gal4* driver and flies *UAS-pallidin-RNAi*^HMS05728^ flies (**A**, n=62-71) or *UAS-pallidin-RNAi*^GD13391^ (**B**, n=44-57) causes reduced sleep time, especially in the first half night (ZT13-18) and long latency. *: p<0.05; **: p<0.005; ***: p<0.0005; Kruskal-Wallis with post hoc comparisons between control and mutant conditions.

**Supplemental Figure S4:**
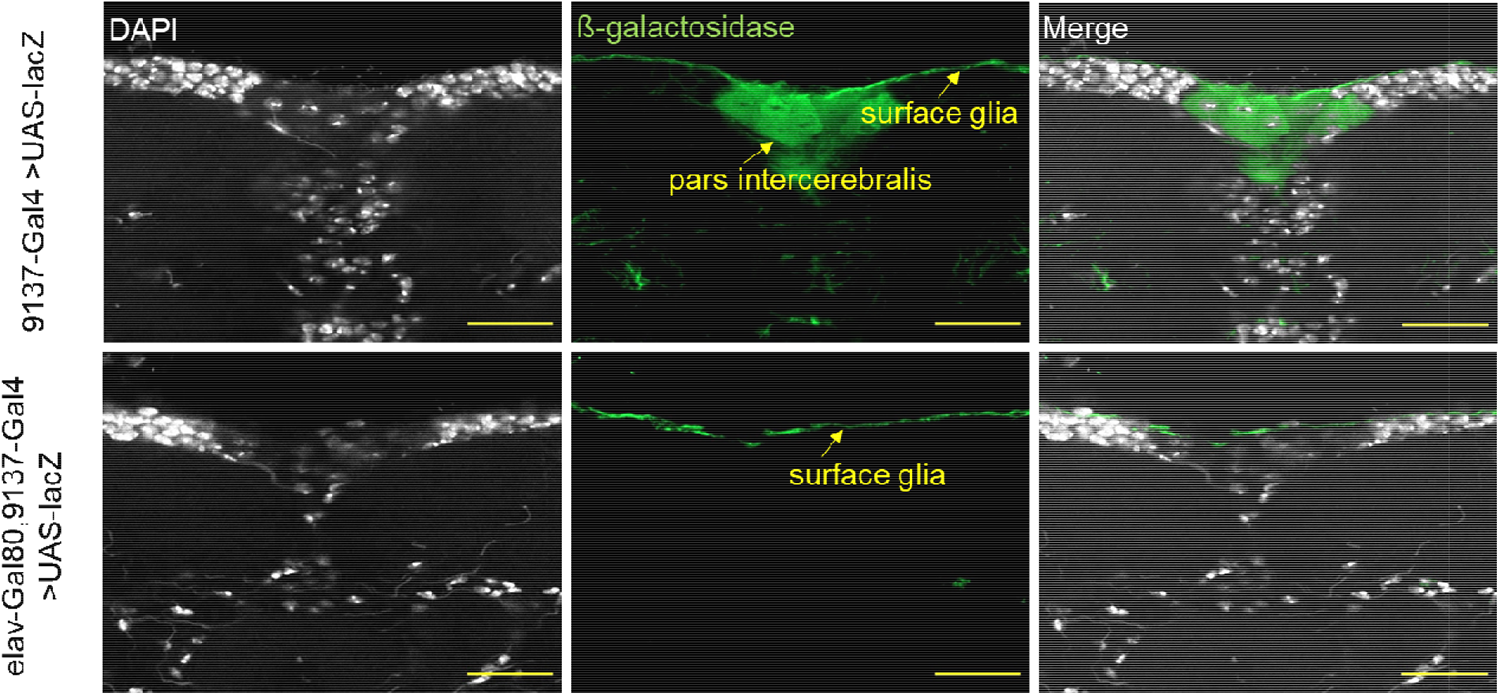
Elav-Gal80 inhibition of 9-137-Gal4 in the pars intercerebralis neurons. Immunofluorescence images of brains with *9-137-Gal4* (top line) or *Elav-Gal80*, *9-137-Gal4* (bottom line) driving a *UAS-lacZ* transgene. *UAS-lacZ* expression is revealed using an anti-ß-galactosidase antibody (green); The ß-galactosidase expression is undetected in the pars intercerebralis neurons when *9-137-Gal4* is combined with *Elav-Gal80*. Single confocal sections. Bar =10 μm, N=4 brains.

**Supplemental Figure S5:**
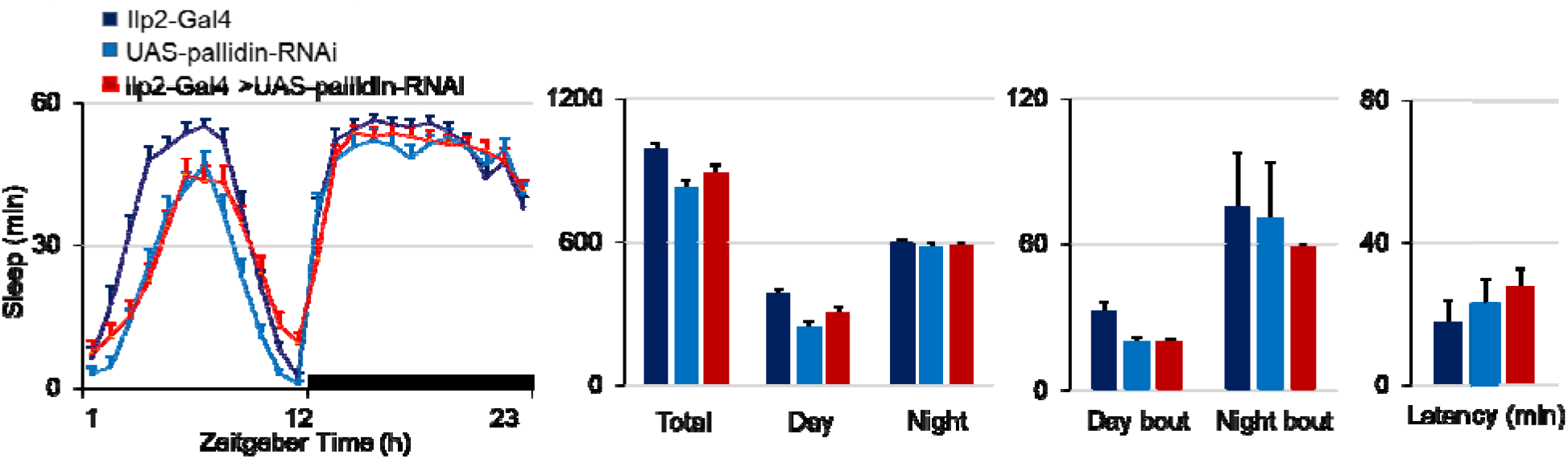
Knockdown of *pallidin* in Ilp2-expressing neurons does not disrupt sleep. 24 hours hourly sleep in Ilp2-Gal4 > *UAS-pallidin-RNAi*^HMS05728^ flies (red) compared with their respective genetic control groups (dark and light blue). Pallidin knockdown flies display similar sleep phenotype to the two controls throughout the day and night, N=30.

**Supplemental Figure S6:**
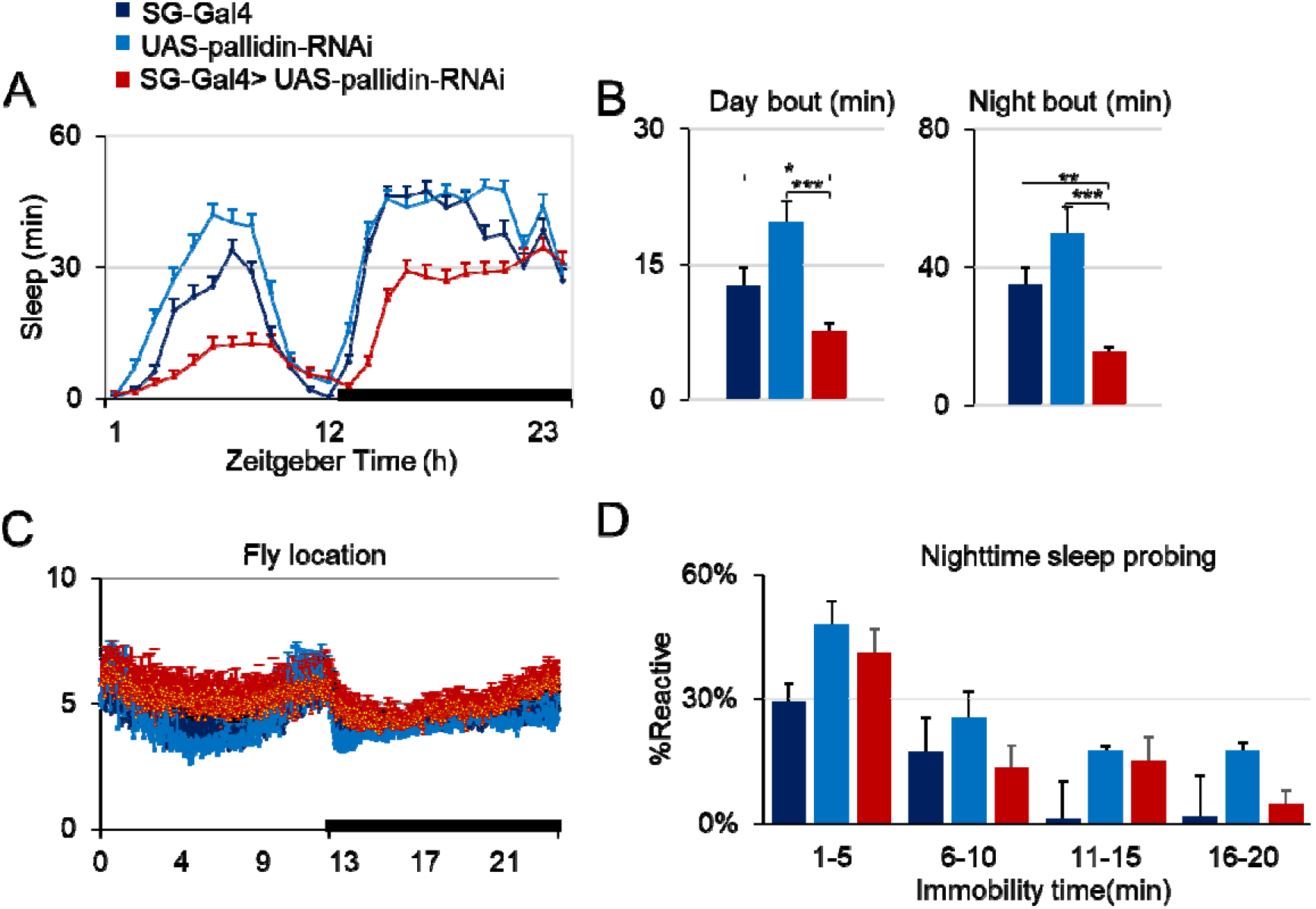
Sleep profiles and sleep intensity in *pallidin* knockdown flies as evaluated with sensory stimulation and videotracking. (A-B). The sleep phenotype of *pallidin* knockdown in surface glia (*elavGal80, 9-137-Gal4* > *UAS-pallidin-RNAi*^HMS05728^) as evaluated with video tracking replicates the results obtained with the infrared beam system. *: P<0.05 **: P<0.005 ***: P<0.0005; Kruskal-Wallis with post hoc comparisons between control and knockdown conditions. N=14-16. (C). Preferential location close to food during sleep was unaffected. 0 is the closest to the food, 10 the farthest; (D).Night sleep probing in *pallidin* knockdown flies and controls. Proportion of flies responding for each immobility bin. Sleep intensity was measured with the DART system: % of flies showing a locomotor response to a 1 sec vibration stimulation, with respect to previous immobility time (0-5min, 6-10min, 11-15min, 16-20min). During the night, knockdown flies and controls showed lower responses above 5 minutes of immobility, the threshold for sleep. This was not observed during the daytime where flies were more responsive to the same stimulation, suggesting lighter sleep, N=45-47.

**Supplemental Figure S7:**
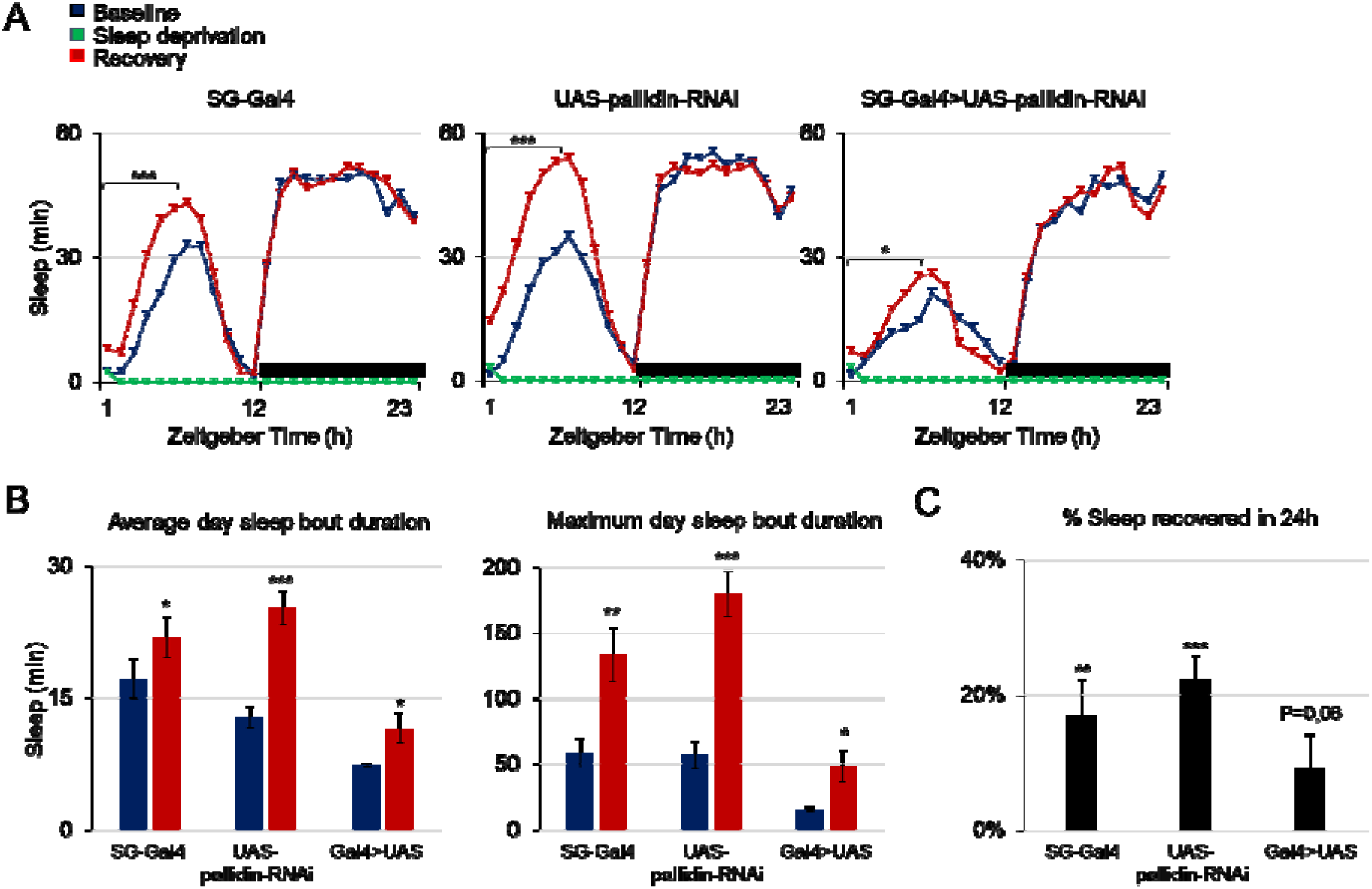
*Pallidin* downregulation modulates the sleep homeostatic response to sleep deprivation. **A**) *pallidin* knockdown (*elavGal80, 9-137-Gal4 > UAS-pallidin-RNAi^HMS05728^*, right panel) and genetic control flies (left and center panel) under baseline condition (blue) were sleep deprived for 24 hours (green) and then allowed to recover for 24h (red). Controls show a clear sleep rebound after sleep deprivation. In the *pallidin* knockdown flies, an increase in sleep is only detected in the first half of the light phase of the recovery day. *: P<0.05; ***: P<0.0005; paired t-test. **B**) Average sleep bout duration (left) and maximum sleep bout duration (right) during the day (light phase) for the baseline day (blue) and for recovery day 1 (red). All groups show a significant increase for these two parameters during recovery. *: P<0.05; **: P<0.005; ***: P<0.0005; paired t-test. **C**) % of sleep recovered for 24h after sleep deprivation, for the different genotypes shown in A. The control groups display a response significantly different from zero while the knockdown are only marginally significant. **: P<0.005; ***: P>0.0005; one sample t-test (N=33-45).

**Supplemental figure S8:**
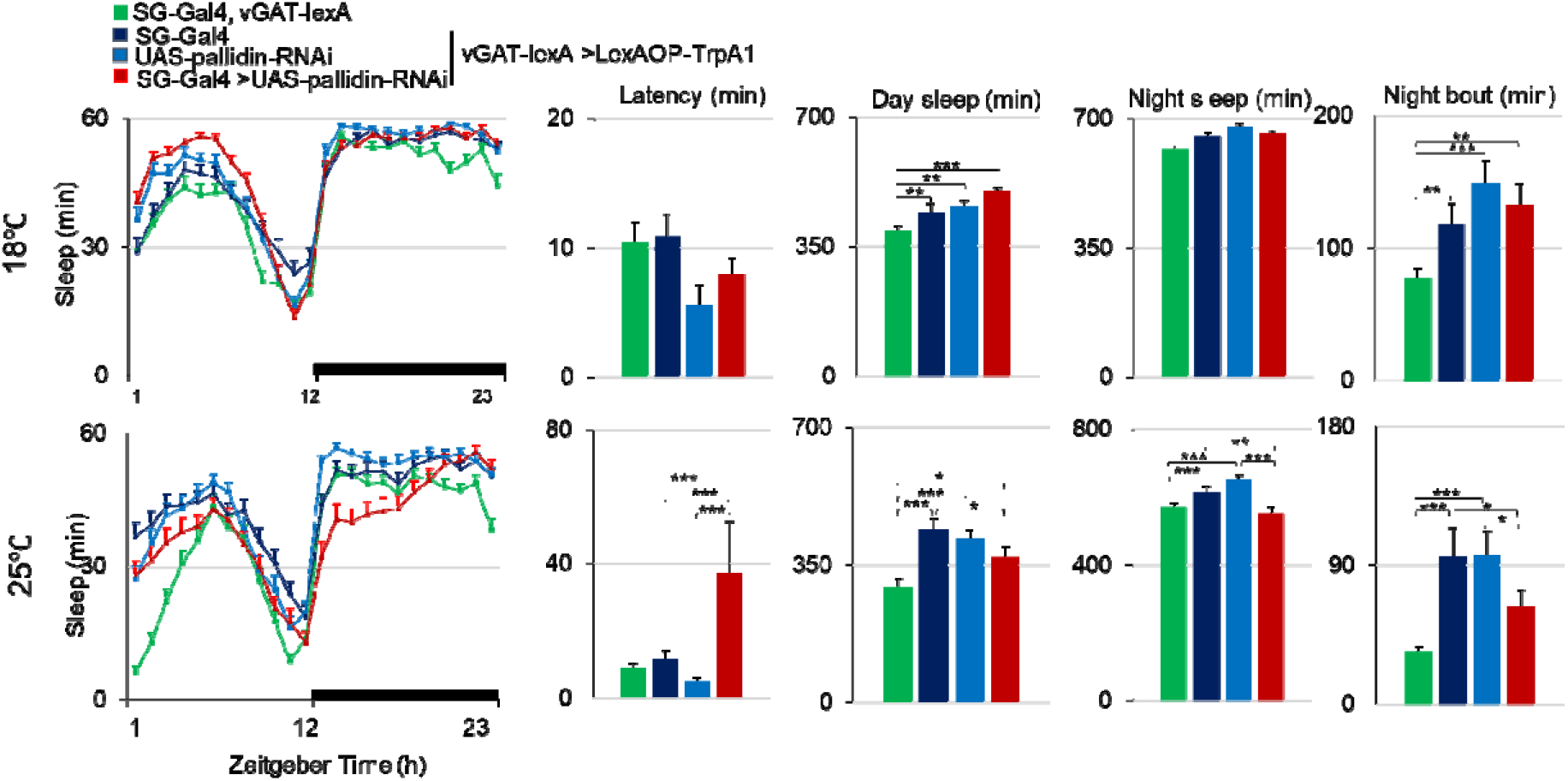
Effect of *pallidin* knockdown in surface glia on GABAergic transmission. Results obtained with *UAS-pallidin-RNAi^iHMS05728^*. Expression of TrpA1 in GABAergic neurons results in higher sleep amounts and longer sleep bout duration at 25°C, compared to non-TrpA1 expressing controls (green), but not at 18°C (upper panel) when the channels are not activated. *Pallidin* knockdown flies (red) did not display higher sleep amounts in response to GABAergic activation, in contrast to their genetic background controls (dark and light blue) (N=40-45). *: p<0.05; **: p<0.005; ***: p<0.0005; Kruskal-Wallis with post hoc comparisons between control and knockdown conditions.

**Supplemental Figure S9:**
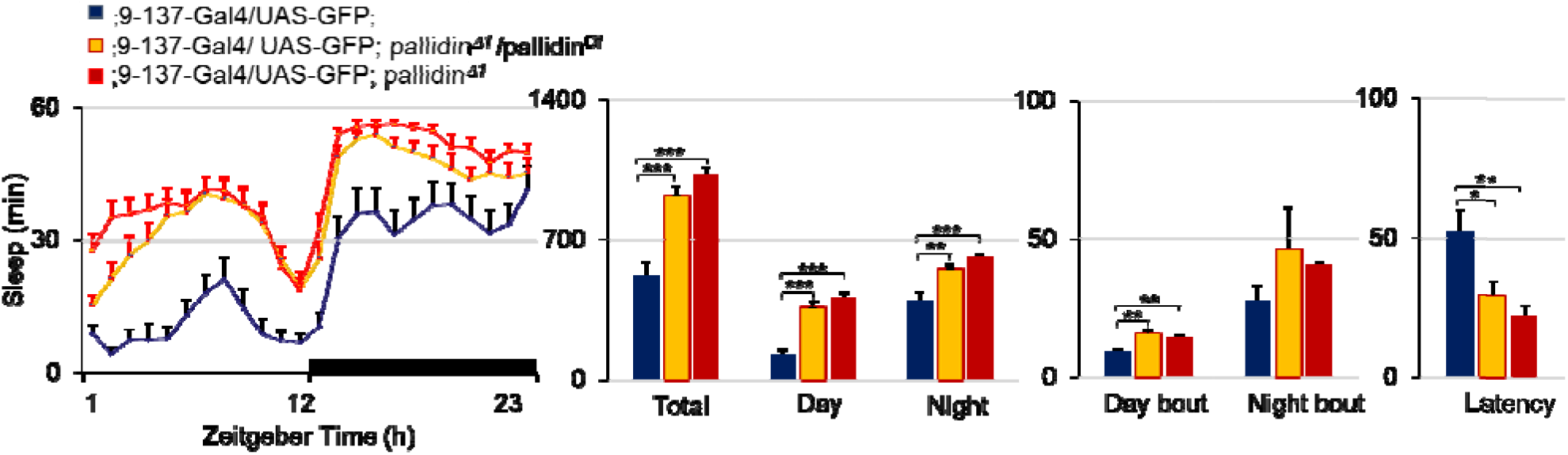
Pallidin*^Δ^*^1^ mutant flies display disrupted sleep. Left, baseline daily sleep and right, sleep architecture parameters of homozygous *pallidin^Δ1^* mutant flies (red) and *pallidin*^Δ1^ / pallidin deficiency (Df(3L)BSC675, orange). Sleep is increased throughout day and night. *: p<0.05; Kruskal-Wallis with post hoc comparisons between control and knockdown conditions, N=17-23.

**Supplemental Figure S10:**
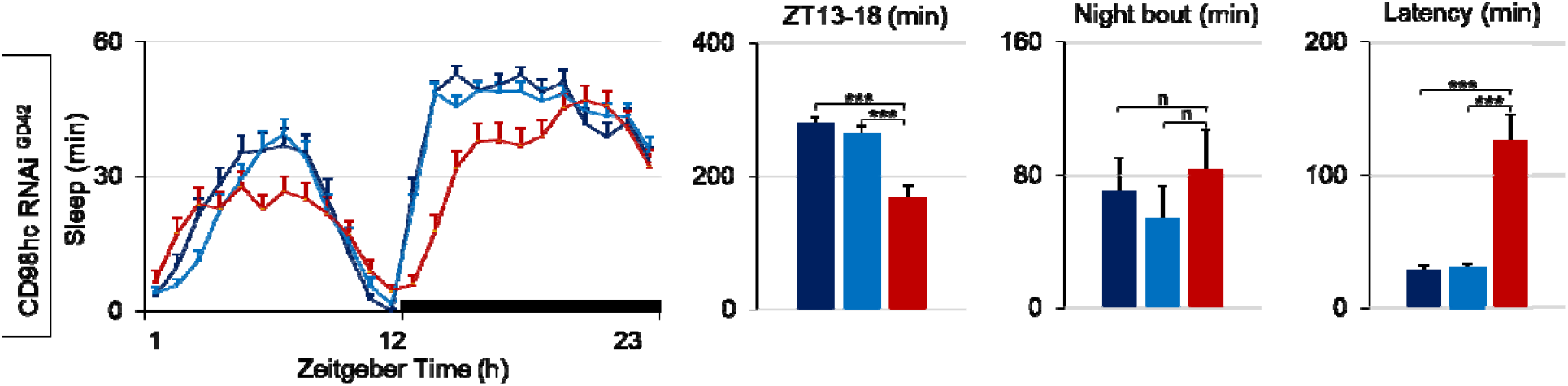
CD98hc downregulation in surface glia results in sleep reduction and long latency. Hourly sleep upon expression of *UAS-CD98hc-RNAi^46622GD^* driven in surface glial cells (*9-137-Gal4* driver) in red, compared to the respective genetic control groups (in blue). Nighttime sleep is reduced, and latency is increased . ***: p<0.0005; Kruskal-Wallis with post hoc comparisons between control and knockdown conditions, N=32-36.

**Supplemental figure S11:**
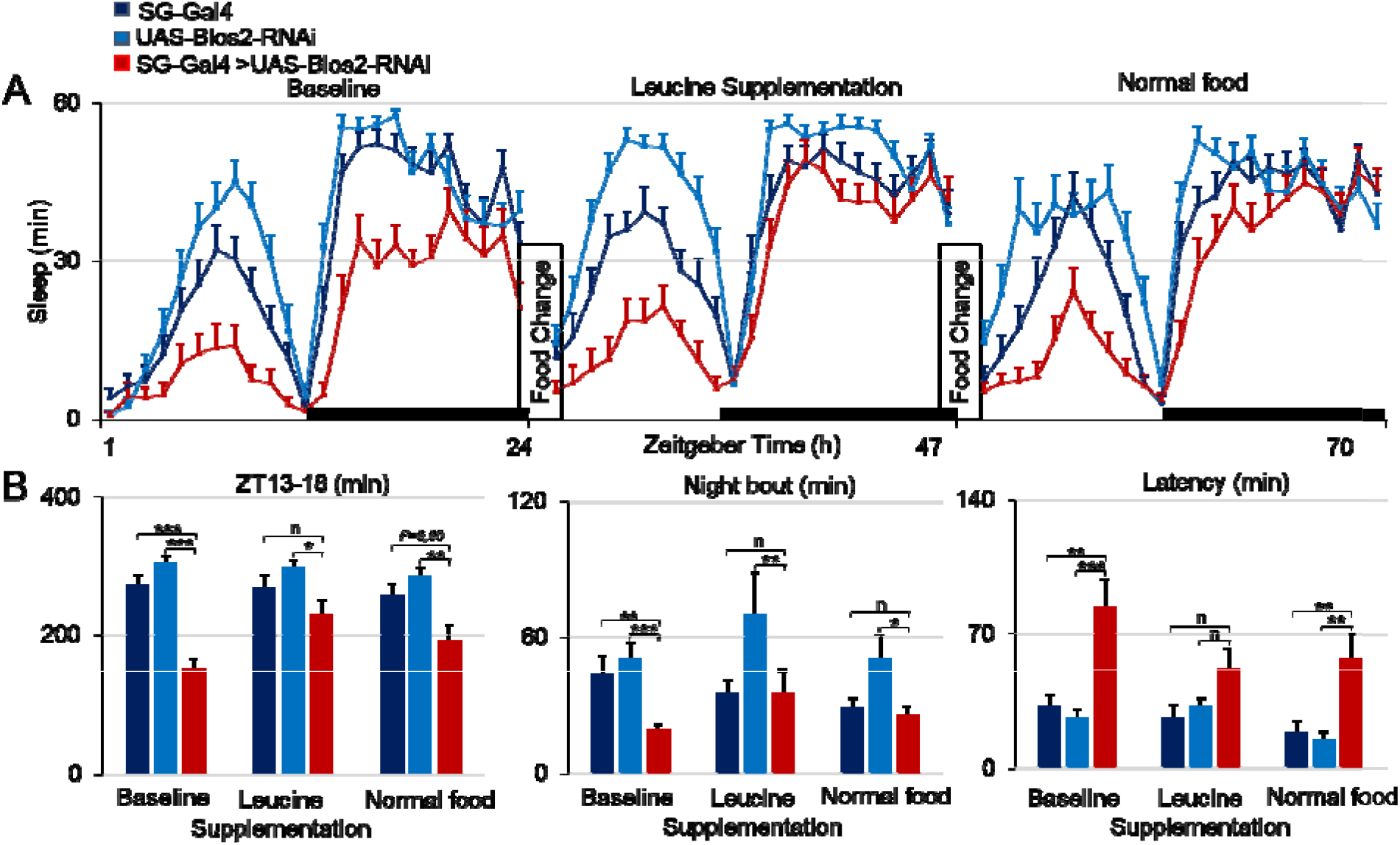
Food supplemented with leucine normalizes the Blos2 downregulation sleep phenotype. Blos2 knockdown flies (*elavGal80,9-137-Gal4>UAS-Blos2-RNAi^GD34355^*) and the genetic background controls fed normal food (baseline) were transferred to fodd supplmemented with leucine for 24h and put back to normal food the following day. Leucine supplementation increased nighttime sleep amounts. *: p<0.05; **: p<0.005; ***: p<0.0005;***; Kruskal-Wallis with post hoc comparisons between control and mutant conditions, N=15-16.

**Supplemental figure S12:**
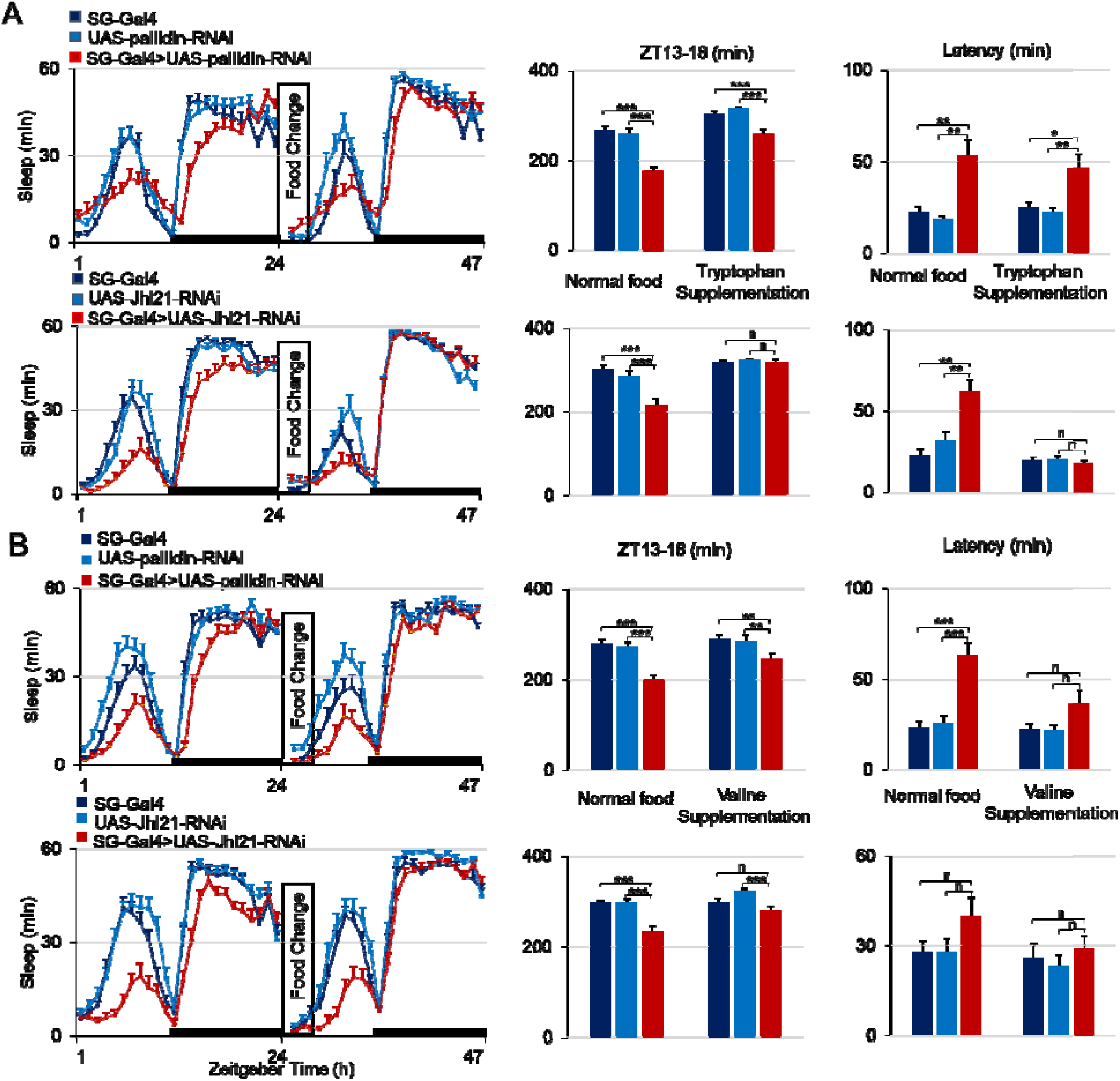
Food supplemented with Tryptophan and Valine partially normalizes the sleep phenotypes of Pallidin and *JhI-21* downregulation in surface glia. (A) *elavGal80, 9-137-Gal4* driving *UAS-pallidin-RNAi^HMS05728^* or *UAS-JhI-21-RNAi^KK112996^* flies (in red) and genetic controls (in blue) were recorded on normal food, then transferred to normal food supplemented with 50 mM Tryptophan for 1 day. (B) The same protocol was applied to valine supplementation for Pallidin or *JhI-21* knockdown flies and their genetic background controls. *: p<0.05; **: p<0.005; ***: p<0.0005;***; Kruskal-Wallis with post hoc comparisons between control and knockdown conditions, N=30-32.

**Supplemental Figure S13:**
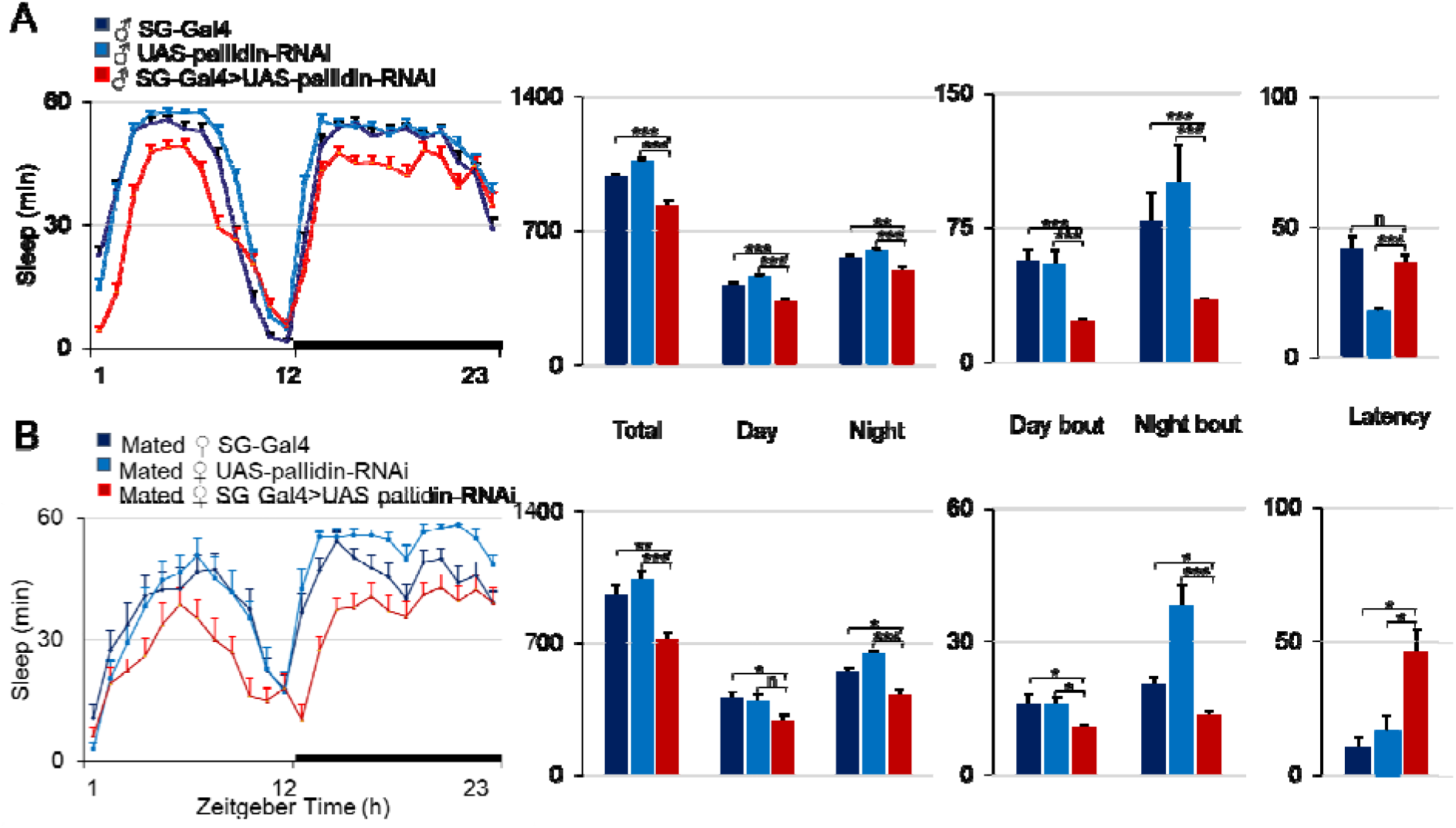
Pallidin function in surface glia is required for sleep independently of mating status and sex. (A) Baseline daily sleep of male flies (*elavGal80, 9-137-Gal4 > UAS-pallidin-RNAi^MS0572^*) (red) and the two genetic controls (dark and light blue) replicate the phenotype observed in females except for the nighttime sleep latency. (B) Baseline daily sleep of mated female flies. Nighttime sleep is reduced, delayed, and fragmented as in non-mated females. *: p<0.05; **: p<0.005; ***: p<0.0005 . Kruskal-Wallis with post hoc comparisons between control and knockdown conditions, N=41-46.

**Supplemental figure S14:**
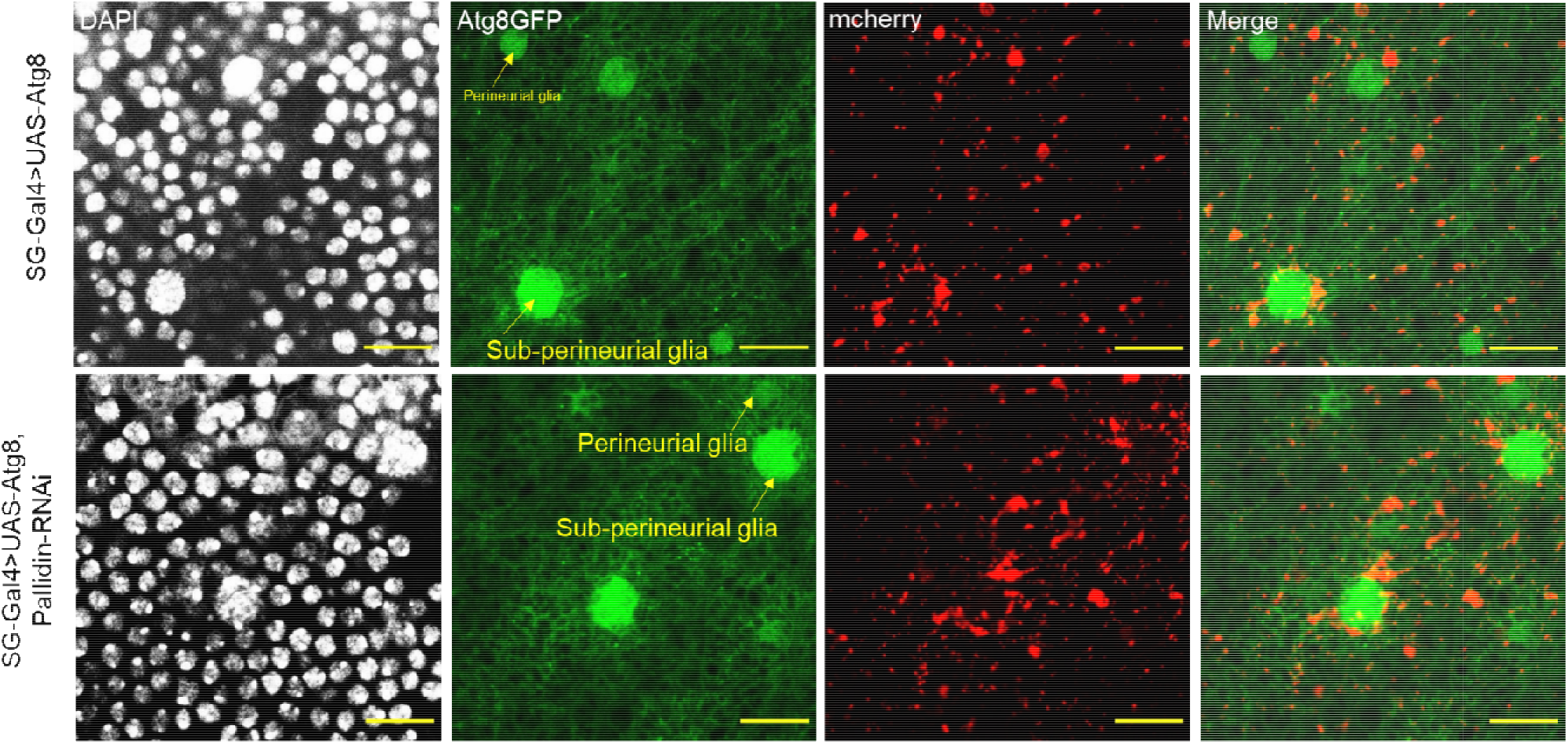
Pallidin down regulation did not result in major changes in autophagy in surface glia. Brains expressing either *UAS-GFP-mcherry-ATG8* alone (controls) or *UAS-GFP-mcherry-ATG8* and *UAS-pallidin-RNAi* driven by *elav-Gal80, 9-137-Gal4*. DAPI (gray), GFP (green) and mcherry (red) fluorescence. Lysosomes labelled by GFP-mcherry-ATG8 were preferentially localized in the perinuclear region of surface glial cells. GFP fluorescence was detected in the nucleus as observed in previous reports (Jacomin et al. Cell reports, 2020). Single confocal section. Pallidin downregulation did not result in obvious changes compared to the control condition. Bar =10μm, N=8.

**Supplemental figure S15:**
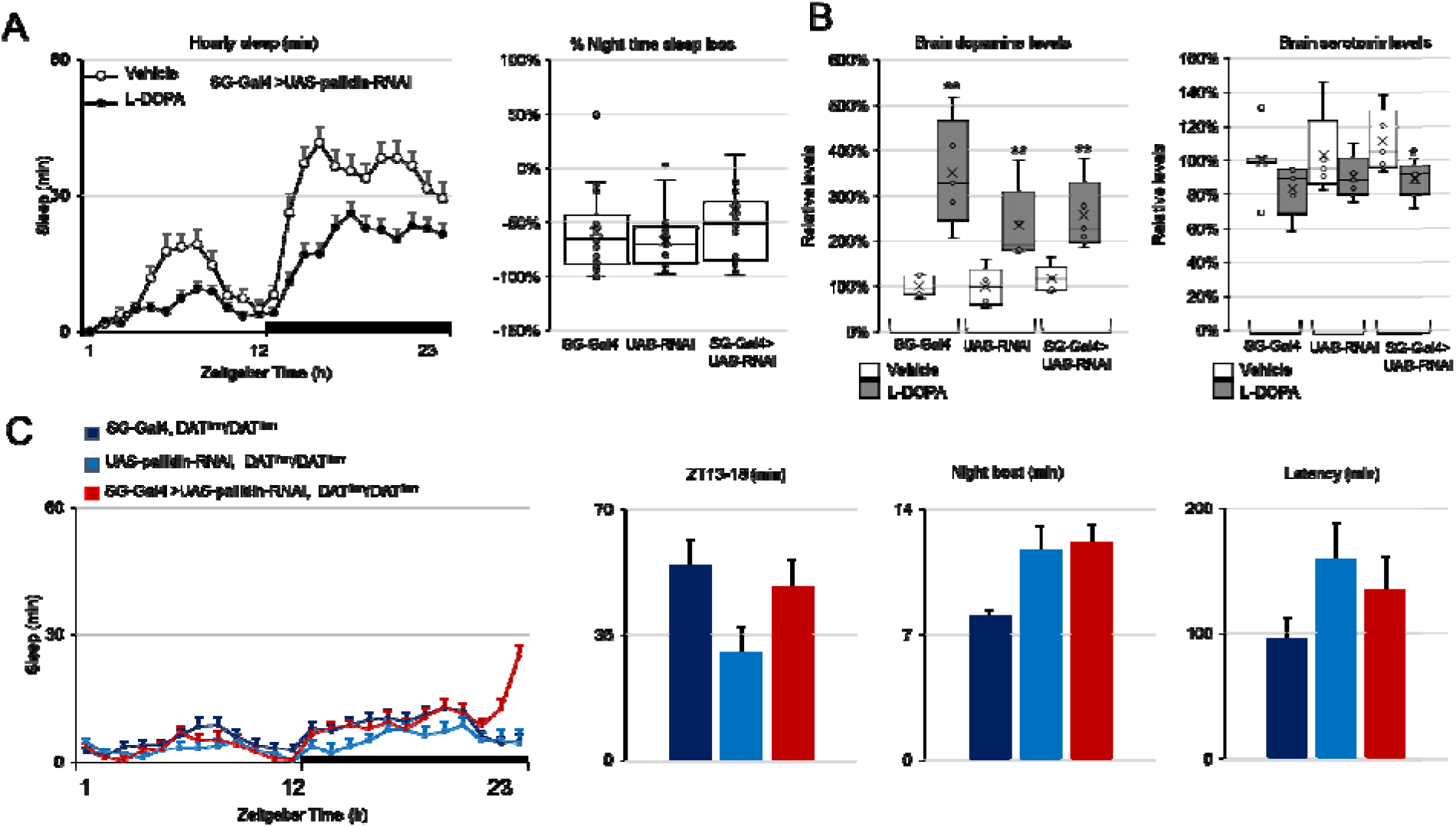
*Pallidin* down-regulation in surface glia failed to increase or attenuate the severity of *fumin*-driven insomnia. A) Hourly sleep in response to L-DOPA feeding. Flies were transferred to L-DOPA or Vehicle at ZT0 (left), the % of night time sleep loss was not significantly different between genotypes (right, whisker box plot). Food intake was similar across genotypes: SG-Gal4: 0.79±0.4; UAS-RNAi: 0.71±0.4; SG-Gal4 >UAS-RNAi: 0.68±0.3 µL/24h N=6) and is not likely to constitute a bias in this experiment. B) Global brain dopamine (left) and serotonin (right) levels relative to the SG-Gal4 control on Vehicle after 24h feeding. Whisker box plots, *: p<0.05; **: p<0.01 Kruskal-Wallis with post hoc comparisons between L-DOPA and Vehicle for each genotype. C) Hourly sleep for knockdown of *pallidin* (*UAS-pallidin-RNAi^GD13391^*) with the *fumin* mutation of the dopamine transporter (DAT) gene in surface glia (in red, N=40) and their controls (in blue, N=41,33). All groups showed similar severe sleep reduction throughout 24h.

## Notes

### Competing Interest Statement

The authors have declared no competing interest.

### Summary of Updates

We have restructured the manuscript, mainly the order and content of the figures, to better highlighting the logical links. A figure showing direct contacts between GABAergic neurons and surface glia is now included. We have added missing details and justifications in the results section and a few more points in the discussion.

